# Developmental regulation of endothelial-to-hematopoietic transition from induced pluripotent stem cells

**DOI:** 10.1101/2024.09.24.612755

**Authors:** Rachel Wellington, Xiaoyi Cheng, Clyde A. Campbell, Cole Trapnell, Raquel Espin-Palazon, Brandon Hadland, Sergei Doulatov

## Abstract

Hematopoietic stem cells (HSCs) arise in embryogenesis from a specialized hemogenic endothelium (HE). In this process, HE cells undergo a unique fate change termed endothelial-to-hematopoietic transition, or EHT. While induced pluripotent stem cells (iPSCs) give rise to HE with robust hemogenic potential, the generation of bona fide HSCs from iPSCs remains a challenge. Here, we map single cell dynamics of EHT during embryoid body differentiation from iPSCs and integrate it with human embryo datasets to identify key transcriptional differences between *in vitro* and *in vivo* cell states. We further map ligand-receptor interactions associated with differential expression of developmental programs in the iPSC system. We found that the expression of endothelial genes was incompletely repressed during iPSC EHT. Elevated FGF signaling by FGF23, an endothelial pathway ligand, was associated with differential gene expression between *in vitro* and *in vivo* EHT. Chemical inhibition of FGF signaling during EHT increased HSPC generation in the zebrafish, while an FGF agonist had the opposite effect. Consistently, chemical inhibition of FGF signaling increased hematopoietic output from iPSCs. In summary, we map the dynamics of EHT from iPSCs at single cell resolution and identify ligand-receptor interactions that can be modulated to improve iPSC differentiation protocols. We show, as proof of principle, that chemical inhibition of FGF signaling during EHT improves hematopoietic output in zebrafish and the iPSC system.

## INTRODUCTION

Hematopoietic stem cells (HSCs) with the potential to sustain the hematopoietic system for life arise in mid-gestation from a specialized population of hemogenic endothelium (HE) lining the dorsal aorta and major embryonic arteries^1–3^. In this process, HE cells undergo endothelial-to-hematopoietic transition (EHT), a unique fate change that involves concordant downregulation of endothelial and induction of hematopoietic expression programs^4,5^. Induced pluripotent stem cells (iPSCs) represent a therapeutically valuable source of human hematopoietic cells^6^. When induced to differentiate into hematopoietic lineages, iPSCs give rise to definitive HE with multilineage hematopoietic potential^7^. However, to date, the generation of bona fide HSCs remains an inefficient process. This broadly suggests that there are differences in hematopoietic ontogeny during iPSC differentiation compared to hematopoiesis in the embryo. While important progress has been made in mapping EHT in human embryogenesis at the single cell level^8–10^, we lack a high-resolution map of EHT during iPSC differentiation in relation to the embryo. Direct comparison between intra-embryonic and iPSC EHT can uncover important differences in developmental gene expression that could be harnessed to improve iPSC differentiation protocols.

In human embryos, the first wave of EHT starts in the yolk sac around Carnegie stage (CS) 7 and involves the production of lineage-restricted erythro-myeloid and multipotent progenitors^2,11^. The generation of HSCs capable of reconstituting hematopoiesis following transplantation begins in the aorta-gonad-mesonephros (AGM) region around CS14^12,13^. Nascent HSCs and progenitors (HSPCs) egress into the bloodstream and colonize the fetal liver around CS16-17 and then bone marrow at birth^12^. It is generally recognized that between CS12-14 (corresponding to E9.5-10.5 in mouse and 24-48 hpf in zebrafish embryogenesis) intra-embryonic HE undergoes developmental maturation to produce HSC-competent HE^8,14^. In the iPSC system, temporal dynamics of EHT have also been well defined^15–17^. Definitive CD34^+^CD43^-^CD73^-^CD184^-^ HE first arises from mesodermal precursors around day 6 of human iPSC differentiation and peaks around day 8^16^. Definitive HE can be further isolated based on expression of CD32 and can undergo robust EHT to give rise to multilineage hematopoietic progenitors^15–17^. Despite the difference in time scales, iPSC-derived hematopoiesis generally recapitulates the stages of embryo hematopoietic ontogeny, critically arterialized endothelium, a precursor to HSC-competent HE^18,19^, and definitive CD32^+^ HE capable of clonal multilineage hematopoiesis^17^.

Since EHT is a transient cell state, there has been considerable interest in capturing this developmental transition using high-depth single cell RNA sequencing (sc-RNA-seq). A number of reports have provided a single cell map of EHT during mouse embryogenesis^20–24^. In human embryos, Zeng et al.^8^ first mapped EHT during early human embryogenesis (CS10-13), a developmental window preceding HSC emergence. Crosse et al.^9^, and more recently Calvanese et al.^10^, have generated single cell maps of EHT during the window of HSC formation (CS14-17) in human embryogenesis. Although several single cell transcriptomic studies have captured iPSC-derived EHT^8,10,15,18,25,26^, we still lack a high-resolution map of stepwise gene expression changes during EHT in iPSC differentiation with direct comparison to hematopoiesis in the embryo. Key regulators of hematopoietic ontogeny including transcription factors (e.g., RUNX1, GATA2, c-MYB), epigenetic regulators (EZH1/2, SUV39), signaling factors (NOTCH, WNT, TGFβ), inflammatory mediators (NFκB, IFN-γ, IL-1β), and biophysical forces (stretch, blood flow) have been identified (reviewed in ^1^). A better understanding of the stage-specific regulation of these factors is essential for recapitulating developmental cues during iPSC differentiation. Direct comparison between carefully staged *in vitro* and *in vivo* cell states spanning hematopoietic ontogeny would facilitate the identification of targetable pathways to improve iPSC-derived hematopoiesis.

To this end, we map EHT dynamics during embryoid body differentiation from iPSCs and identify key transcriptional differences between *in vitro* and *in vivo* cell states. By mapping ligand-receptor interactions associated with the differential expression of developmental programs, we identify elevated FGF signaling as a barrier to hematopoietic specification. These data identify a panel of ligands that may be modulated to improve iPSC differentiation protocols.

## RESULTS

### Single-cell temporal profiling of hematopoietic ontogeny *in vitro*

The endothelial-to-hematopoietic transition (EHT) is a key event in hematopoietic ontogeny; however, the dynamics of EHT during iPSC differentiation have not been deeply profiled in direct comparison to the embryo. To capture the dynamic transcriptional landscape of EHT, we first performed an unbiased single cell transcriptional profiling of embryoid bodies (EBs) cultured under hematopoietic differentiation conditions. We collected nuclei from whole EBs on days 7, 8, 11, 14, 18 and 21 of differentiation and performed well-based single cell combinatorial indexing RNA-seq (sci-RNA-seq) enabling the capture of a large number of cells for assessment of hematopoietic development (**Figure 1A**). After standard filtration steps (see **Methods**), sequencing data for a total of 166,085 cells across all days of differentiation were acquired (**Figure 1B**) (see Code and Data Availability in **Methods**). Cell populations were identified based on the top relative expression score of previously identified CS12-16 whole embryo cell type gene marker sets^27^ for each individual cell cluster (**Figure 1B, Figure S1A-D**; **Table S1**). We further refined this annotation using extended cell type marker scores (**Figure S1D**). This analysis confirmed that EBs are comprised of cell types from all germ layers: mesoderm (somite, blood, fibroblast, lateral plate mesoderm), endoderm, and ectoderm (neural, Schwann, epithelium). Hematopoietic cells were further characterized based on the expression of *RUNX1*, *SPN* (CD43), and *PTPRC* (CD45), while precursor endothelial populations were identified by the expression of *CDH5*, *KDR*, *TIE1*, and *CD34* (excluding *RUNX1/SPN/PTPRC* hematopoietic cells) (**Figure 1C-D, Figure S1E-F**; **Table S1**). Hematopoietic and endothelial populations were connected by a “bridge”, representing HE cells undergoing EHT to acquire hematopoietic cell fate prior to differentiating into various hematopoietic lineages. Endothelial cells proximal to the bridge acquired arterial markers, such as *DLL4*, consistent with arterial endothelial (AE) cells as the precursor of HE (**Figure 1C**). Notably, EHT was robust in early ontogeny peaking around day 8 and declining thereafter, with fewer cells transiting at day 11, and EHT virtually ceasing by day 18 of differentiation (**Figure 1D, Figure S1F**). At the same time, endothelial and hematopoietic populations became more separated over time, indicating their progressive transcriptional divergence later in ontogeny (**Figure 1D, Figure S1F**). These results show that, akin to embryogenesis, EHT is a transient cell state during iPSC differentiation with the peak in our system around day 8, consistent with the known timing of EHT across various differentiation protocols^16,19,28–31^.

**Figure 1.**
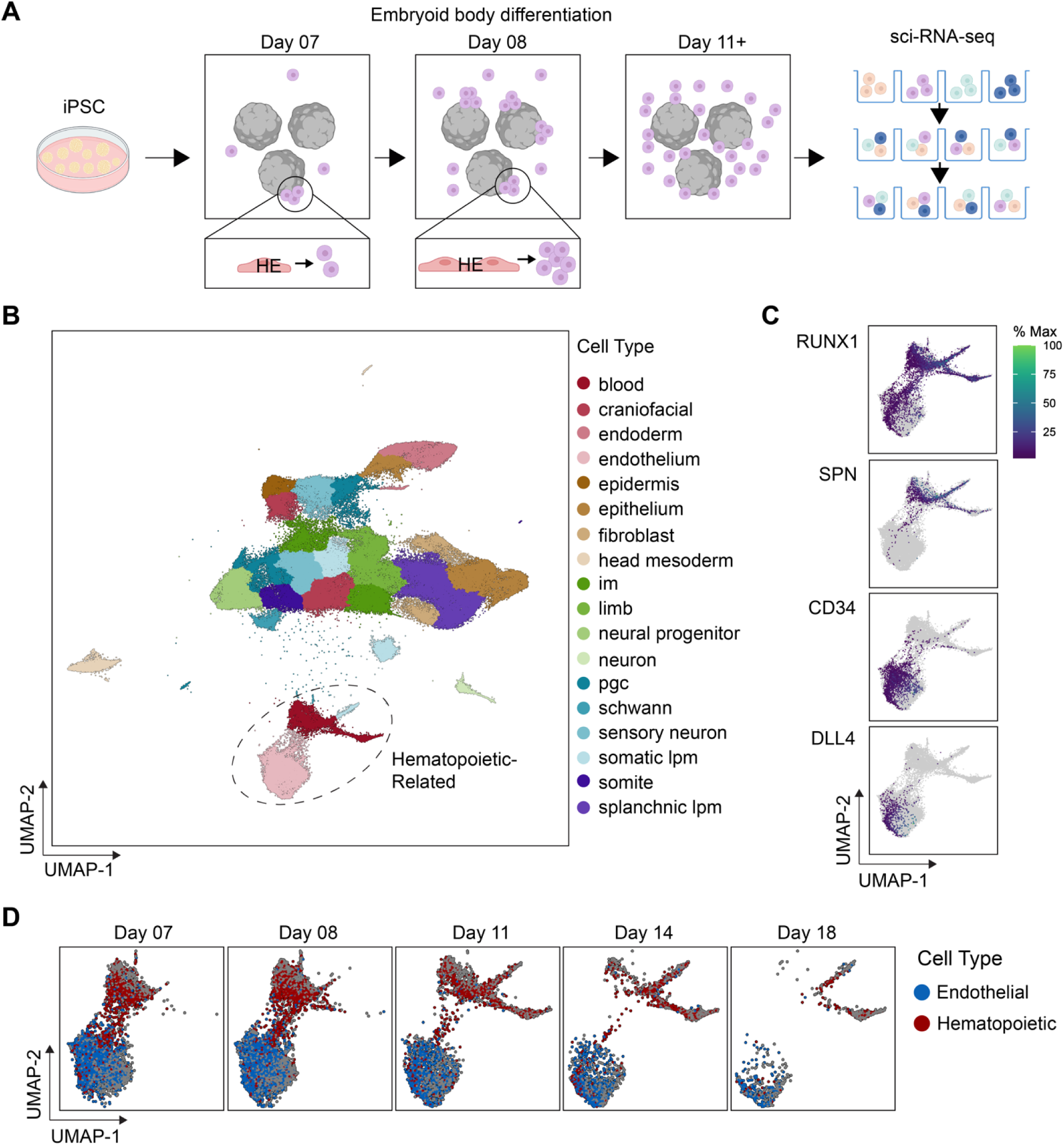
Single cell temporal profiling of hematopoietic ontogeny *in vitro*. (**A**) Experimental scheme of single cell profiling during iPSC differentiation into HSPCs. Whole EBs were dissociated on days 7, 8, 11, 14, 18 and 21 of EB differentiation and nuclei profiled by sci-RNA-seq. (**B**) UMAP dimensional reduction of combined sci-RNA-seq data from days 7, 8, 11, 14, 18 and 21 days of EB differentiation. Cell types are labeled based on the maximal scores of gene signatures from Xu et al.^27^ shown in Figure S1A. The dashed circle contains the hematopoietic and endothelial populations. (**C**) Expression of *RUNX1*, *CD34*, *SPN* (CD43), and *DLL4* transcripts in the hematopoietic and endothelial populations. (**D**) Hematopoietic (red) and endothelial (blue) populations shown by day of differentiation showing the temporal dynamics of hematopoiesis. Cell types were identified as hematopoietic (*RUNX1*, *SPN*, *PTPRC* aggregate gene score) or endothelial (*CDH5*, *KDR*, *TIE1*, *CD34*, and zero hematopoietic aggregate gene score). Panel (A) was created with BioRender.com.

### Developmental dynamics of EHT *in vitro* and *in vivo*

To generate a high-resolution map of EHT from iPSCs, we next focused on the peak of EHT at day 8 of EB differentiation. To enrich for HE cells, we purified endothelial and hematopoietic cells positive for CD34 or CD43/CD45 and performed 10X single cell RNA-seq (sc-RNA-seq) (**Figure 2A, Figure S2A-B**; **Table S2**). As expected, almost all cells in our analysis were endothelial (*CD34*, *SPN*^neg^) expressing arterial marker *DLL4,* or hematopoietic (*RUNX1*, *SPN^+^*) (**Figure 2B**). We captured a robust population of HE cells (*RUNX1*, *CD34*, *DLL4*, *SPN*^neg^) actively undergoing EHT (**Figure 2B-D**). We next identified endothelial and hematopoietic subtypes based on published marker sets^8–10^ and assigned supervised cell type annotations to clusters identified in UMAP space using Monocle 3 (**Figure 2E-G**; **Table S2**). This analysis revealed the following conclusions. First, endothelial cells expressing *DLL4* were proximal to HE. As expected, cluster 27 on the endothelial “side” of the bridge was identified as early HE (**Figure 2C,D**). Interestingly, cluster 26a was enriched for both an HE and HSC-like signature (**Figure 2E,G**), likely representing more mature HE^10^. By contrast, a second trajectory (cluster 26b) did not exhibit the HE or HSC signature, but instead expressed a strong erythrocyte signature, likely representing erythro-myeloid precursors (**Figure 2C-E**). Both trajectories were linked to hematopoietic precursors and lineages, including erythroid and myeloid. These data capture the transcriptome changes of maturing HE undergoing EHT, as well as the emergence of more primitive, early erythroid precursors.

**Figure 2.**
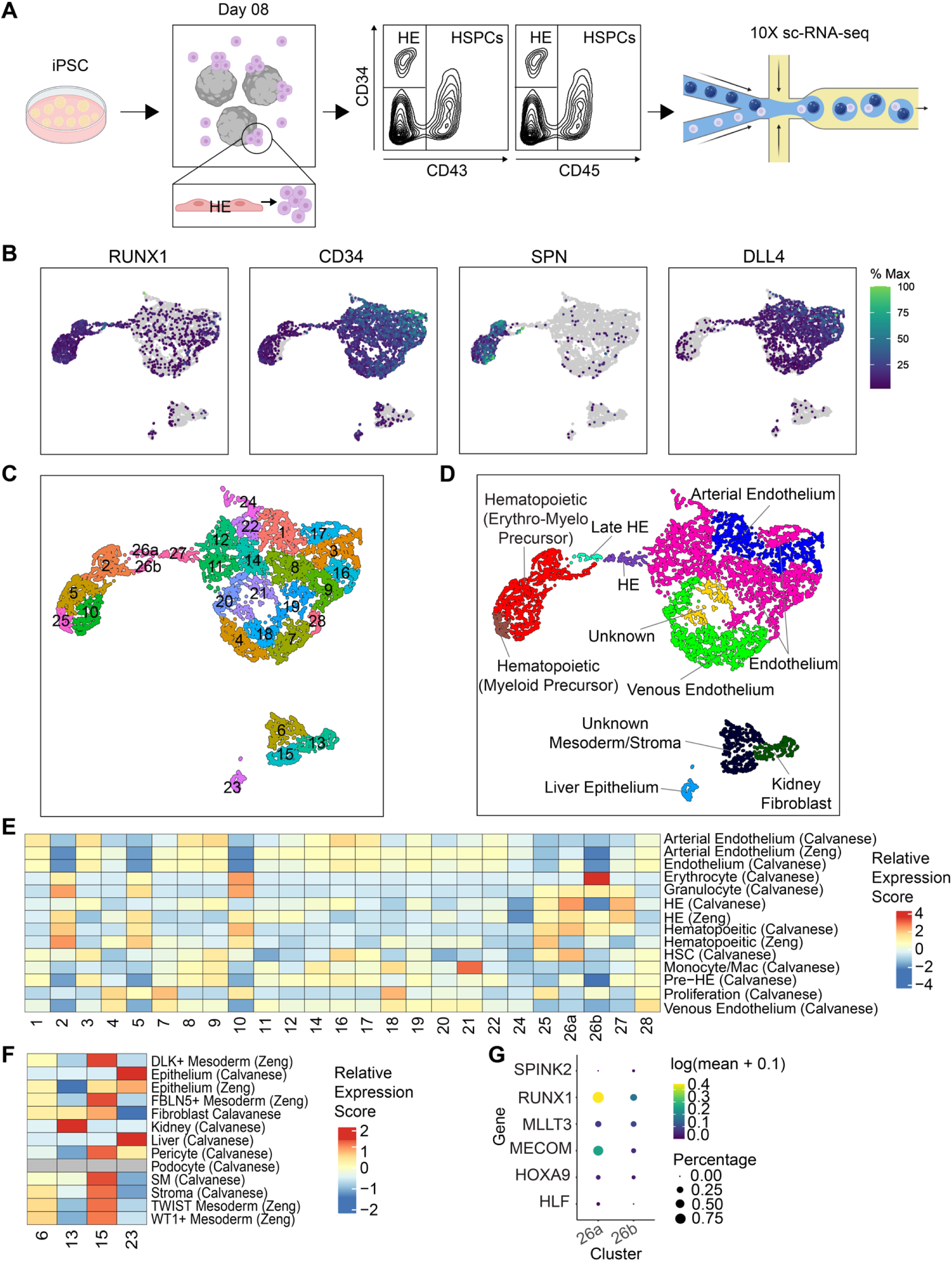
Developmental dynamics of EHT during iPSC differentiation. (**A**) Experimental scheme of 10X single cell RNA-sequencing of EHT on day 8 of differentiation. iPSCs were differentiated as EBs into HSPCs, dissociated on day 8 and hematopoietic and endothelial populations were flow sorted for profiling by 10X 3’ sc-RNA-seq based on the expression of CD34, CD43, and CD45, as shown. (**B**) Expression of *RUNX1*, *CD34*, *SPN* (CD43), and *DLL4* transcripts. (**C**) UMAP showing unsupervised clustering. (**D**) Cell type classification with cell types identified by top gene signatures in E-F. (**E,F**) Heatmap of relative expression (aggregate gene scores in a scaled matrix) of published gene signatures for each cluster in C (Table S2). (**G**) Expression of HSC genes from Calvanese et al.^10^ in HE clusters 26a and 26b. Panel (A) was created with BioRender.com.

To uncover molecular pathways essential for HSC generation, we next sought to compare the dynamics of EHT from iPSCs to HSC-competent human embryos. To this end, we first integrated our EB day 8 sc-RNA-seq data with published human embryo sc-RNA-seq datasets using Seurat canonical correlation analysis^32^ (see **Methods**). The published human embryo datasets include CS10-13 preceding HSC formation from Zeng et al^8^, CS16 from Crosse et al.^9^, and CS14-17 from Calvanese et al.^10^, the latter two studies spanning active HSC generation. iPSC-derived and human embryo datasets overlapped in common UMAP space (**Figure 3A-B**; **Table S3**). We identified HE undergoing EHT (*RUNX1^+^*, *CD34^+^*, *DLL4^+^*, *SPN*^neg^) at CS10-CS13 and CS14 time points, with increasing separation of the endothelial (*CD34^+^*, *CD43*^neg^) and hematopoietic (*RUNX1^+^*, *SPN*^+^) components at CS15 and CS17 (**Figure 3A-C**), resembling what we observed in our initial *in vitro* time course (**Figure 1D**). As in the iPSC data, *DLL4*^+^ AE (cluster 36) preceded the *RUNX1*^+^ HE “bridge” (cluster 75) giving rise to all hematopoietic lineages (**Figure 3B-C**). We further rigorously validated that developmentally equivalent populations were correctly integrated by transferring cell labels from our day 8 EB data, as well as each individual published dataset onto the common UMAP space (**Figure 3D-E**, **Figure S3A-E**; **Table S3**). Our iPSC-derived and integrated datasets can be explored interactively through an online interface (see **Code and Data Availability**). These data suggest that iPSC-derived HE undergoes similar developmental transitions to those in human embryo AGM enabling their direct molecular comparison.

**Figure 3.**
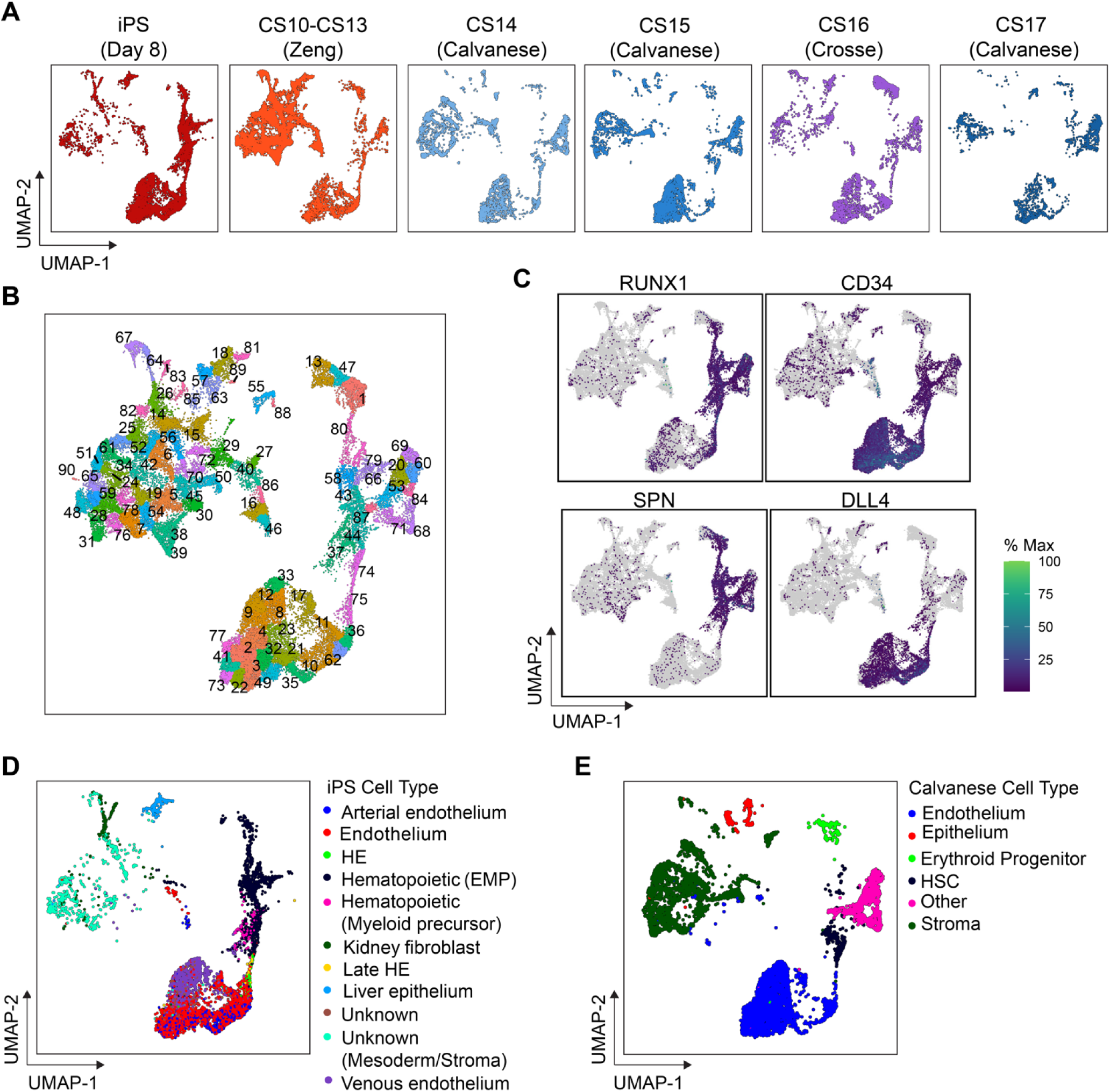
Developmental dynamics of EHT *in vitro* compared to *in vivo*. (**A**) Integration of day 8 iPSC-derived EB and human embryo 10X sc-RNA-seq data from Zeng et al.^8^, Crosse et al.^9^ and Calvanese et al.^10^ into the same UMAP dimensionally-reduced space using Seurat canonical correlation analysis, with individual datasets separated post-integration. (**B**) Integrated UMAP with unsupervised clustering in Monocle 3. (**C**) Expression of *RUNX1*, *CD34*, *SPN* (CD43), and *DLL4* transcripts. (**D,E**) Transfer of cell type labels from the day 8 EB dataset (D) and Calvanese et al.^10^ (E) onto the integrated space for verification of proper integration.

### Comparative molecular analysis of EHT *in vitro* and *in vivo*

To uncover molecular pathways essential for HSC generation, we next compared transcriptome changes during EHT between iPSC and HSC-competent human embryos (CS14-17). We focused on the AE preceding EHT (cluster 36), HE undergoing EHT (cluster 75), and emergent HSPCs exiting EHT (cluster 74) in our integrated dataset (**Figure 3B**). We first identified differentially expressed genes (DEGs) during the entire EHT (comprising HE and emergent HSPCs) between iPSC and HSC-competent human embryo using a mixed-negative binomial model accounting for batch effects in Monocle 3 (**Figure S4A**; **Table S4A**). DEGs downregulated in the iPSC EHT were enriched for gene ontology (GO) terms related to cytokine signaling and hemopoiesis (**Figure S4B,C**; **Table S4B**). Consistent with HSC-incompetent EHT, significantly downregulated genes in iPSC-derived cells included transcription factors essential for HSC and lymphoid fate specification, including *HOXA5/7/10*, *HLF*, *MLLT3*, *SPINK2*, *SPI1*, *GFI1*, *GATA2*, *TCF7*, and *BCL11A*, and signaling receptors including *CD37*, *CD40*, *CD44*, *CD52*, *CD68*, *CD74*, *CSF1R*, *IL6R*, *IL3RA*, and *IL11RA* (**Table S4A**). Since most of these transcriptional changes likely reflect the HSPC output of EHT, we next focused on HE that has not yet acquired hematopoietic identity (**Figure 4A**; **Table S4A**). Genes downregulated in iPSC HE were enriched for GO terms related to cytokine signaling, hematopoiesis, and HSC homeostasis (**Figure 4B,C**; **Table S4B**). Key downregulated genes in iPSC HE included transcriptional regulators *SOX17*, *GATA2*, *REL*, *ID2*, and *NKX2-3* and signaling factors *ALDH1A1* (retinoic acid signaling), *CD74*, *IL11RA*, and *IL3RA* (**Table S4A**). Upregulated genes in iPSC HE were enriched for GO terms related to mitochondrion organization (**Figure 4B**; **Table S4B**), and surprisingly included endothelial genes *KDR* and *FLT1*. Lastly, genes downregulated in iPSC-derived early HSPCs were also enriched for GO terms related to cytokine signaling and hemopoiesis (**Figure S4D-F**; **Table S4B**).

**Figure 4.**
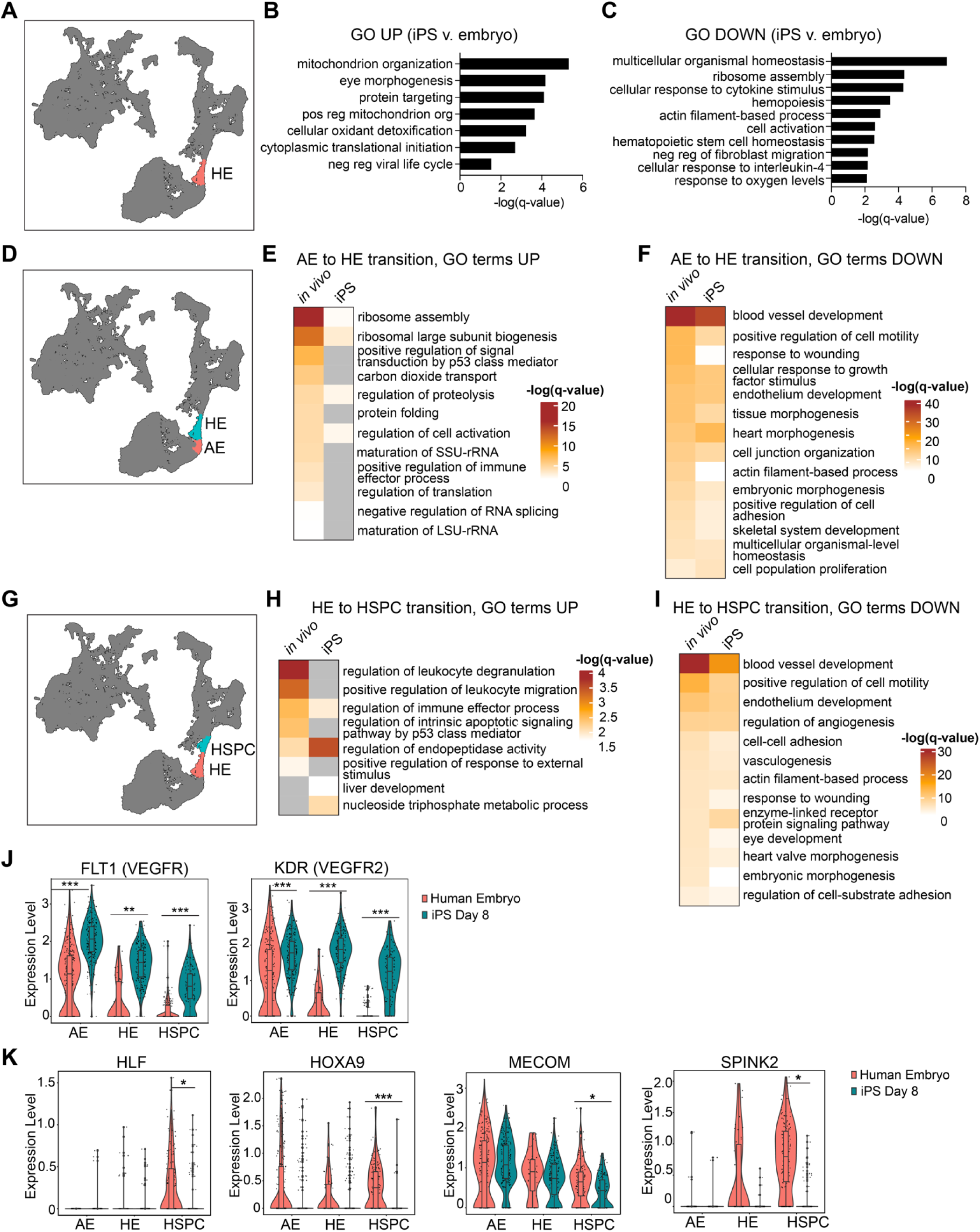
Comparative molecular analysis of EHT *in vitro* and *in vivo*. (**A**) HE population (cluster 75 in integrated dataset) used for differential gene expression analysis. (**B,C**) Gene ontology (GO) terms significantly upregulated (B) or downregulated (C) in iPSC-derived vs. HSC-competent human embryo HE (Metascape, q < 0.05; top 10 terms). (**D**) Arterial endothelium (AE; cluster 36, red) and HE (cluster 75, blue) populations used for differential expression analysis. (**E,F**) GO terms significantly upregulated (E) or downregulated (F) during AE-to-HE transition (Metascape batch mode, q < 0.05). (**G**) HE (cluster 75, red) and hematopoietic stem and progenitor cell (HSPC; cluster 74, blue) populations used for differential expression analysis. (**H,I**) GO terms significantly upregulated (H) or downregulated (I) during HE-to-HSPC transition (Metascape batch mode, q < 0.05). (**J**) Expression of representative endothelial identity genes *FLT1* and *KDR* in HSC-competent human embryos (CS14-17; red) and day 8 iPS (blue) AE (cluster 36), HE (cluster 75) and HSPCs (cluster 74). (**K**) Expression of HSC genes *HLF*, *HOXA9*, *MECOM*, and *SPINK2* in HSC-competent human embryos (CS14-17; red) and day 8 iPS (blue) AE (cluster 36), HE (cluster 75) and HSPCs (cluster 74). For (J, K) *q < 0.05, **q < 0.005, and ***q < 0.0005 using a regression fit to a mixed-negative binomial model accounting for batch effects.

We next tracked gene expression dynamics along hematopoietic ontogeny independently in the iPSC system and human embryos. To this end, we identified DEGs in the transition from AE to HE, and the subsequent transition from HE to HSPC, either *in vitro* or *in vivo* (**Table S4A**). In the AE to HE transition, GO terms related to ribosome biogenesis, cell activation, splicing, and translation, were significantly induced in the embryo (**Figure 4D,E, Figure S4G**; **Table S4B**), reflecting a more activated cell state of HE. At the same time, GO terms related to blood vessel development, cell motility, endothelial development, and cell junctions, were downregulated in embryo HE (**Figure 4F, Figure S4H**; **Table S4B**), reflecting a gradual loss of endothelial cell identity. These downregulated GO terms were similarly downregulated in the AE to HE transition in iPSCs (**Figure 4F, Figure S4H**; **Table S4B**), suggesting that the loss of endothelial cell identity occurs appropriately at this stage. In the subsequent developmental transition from HE to HSPC, GO terms related to leukocyte biology and immune process were upregulated in the embryo (**Figure 4G,H, Figure S4I**; **Table S4B**), while terms related to blood vessel development, cell motility, and endothelium development were downregulated (**Figure 4I**; **Table S4B**), reflecting acquisition of hematopoietic identity and loss of endothelial identity. Interestingly, endothelial cell identity was less strongly repressed in the HE to HSPC transition in iPSCs than the embryo (**Figure 4I, Figure S4J**). Consequently, iPSC-derived HE and HSPCs maintained an elevated expression of endothelial-related genes (**Figure S4J**). Among these, key endothelial cell fate genes *FLT1* (VEGFR1) and *KDR* (VEGFR2) were appropriately repressed in embryo-derived HSPCs but continued to be expressed in iPSC-derived HSPCs at significantly higher levels (**Figure 4J**). By contrast, iPSC-derived HSPCs failed to upregulate key HSC genes (**Figure 4K**). Taken together these data suggest that iPSC-derived EHT is characterized by lower expression of HSC-specifying transcription and signaling factors and a surprising failure to properly suppress endothelial cell gene expression.

### A map of ligand-receptor interactions to inform iPSC differentiation

Since iPSC-derived EHT was characterized by misexpression of signaling factors, we reasoned that the hemogenic potential can be augmented by modulation of specific ligand-receptor signaling pathways. To identify candidate signaling pathways, we connected upstream signals with differential gene expression in iPSC-derived EHT using the NicheNetR package^33^. We applied NicheNetR to our integrated dataset and identified ligand-receptor pairs predicted to account for differential gene expression between EHT in iPSCs and HSC-competent EHT in human embryos. We required that receptors be expressed on HE cells, without defining the ligand source to allow for ligands produced by other sources than the cells in our dataset (e.g., hormones or cell culture media components). Top ligand-receptor interactions predicted to signal upstream of genes differentially expressed between *in vitro* and *in vivo* HE included signaling pathways previously implicated in regulating EHT, such as TNF, IFN-γ, TGFα, EDN1, and HAS2 signaling via CD44^1,34^ (**Figure 5A**, **Figure S5A**; **Table S5**). Extending the analysis to the entire EHT population (HE and HSPC) identified TGFβ signaling, HAS2 signaling, and NOTCH signaling pathways, which are known regulators of EHT^1^ (**Figure 5B**, **Figure S5B**; **Table S5)**. Ranking the pathways by the activity score and number of target genes identified TNF, ADAM17, CAMP, IL-13, HAS2, IFNG, TGFA, FGF23, EDN1, and AGT signaling as the top modulable pathways for iPSC-derived HE (**Figure 5C**, left). Extending this analysis to the entire EHT population identified ADAM17, DLL1/DLL4/NOTCH, HAS2, SECTM1, NOV, IL-13, TGFB3, and IL-25 as the top modulable pathways (**Figure 5C**, right). Many of these pathways have not been previously implicated in hematopoiesis or hematopoietic development. These data uncover the signaling pathways that may be mis-regulated in iPSC-derived EHT resulting in suboptimal hematopoietic output and serve as a resource to improve iPSC differentiation protocols.

**Figure 5.**
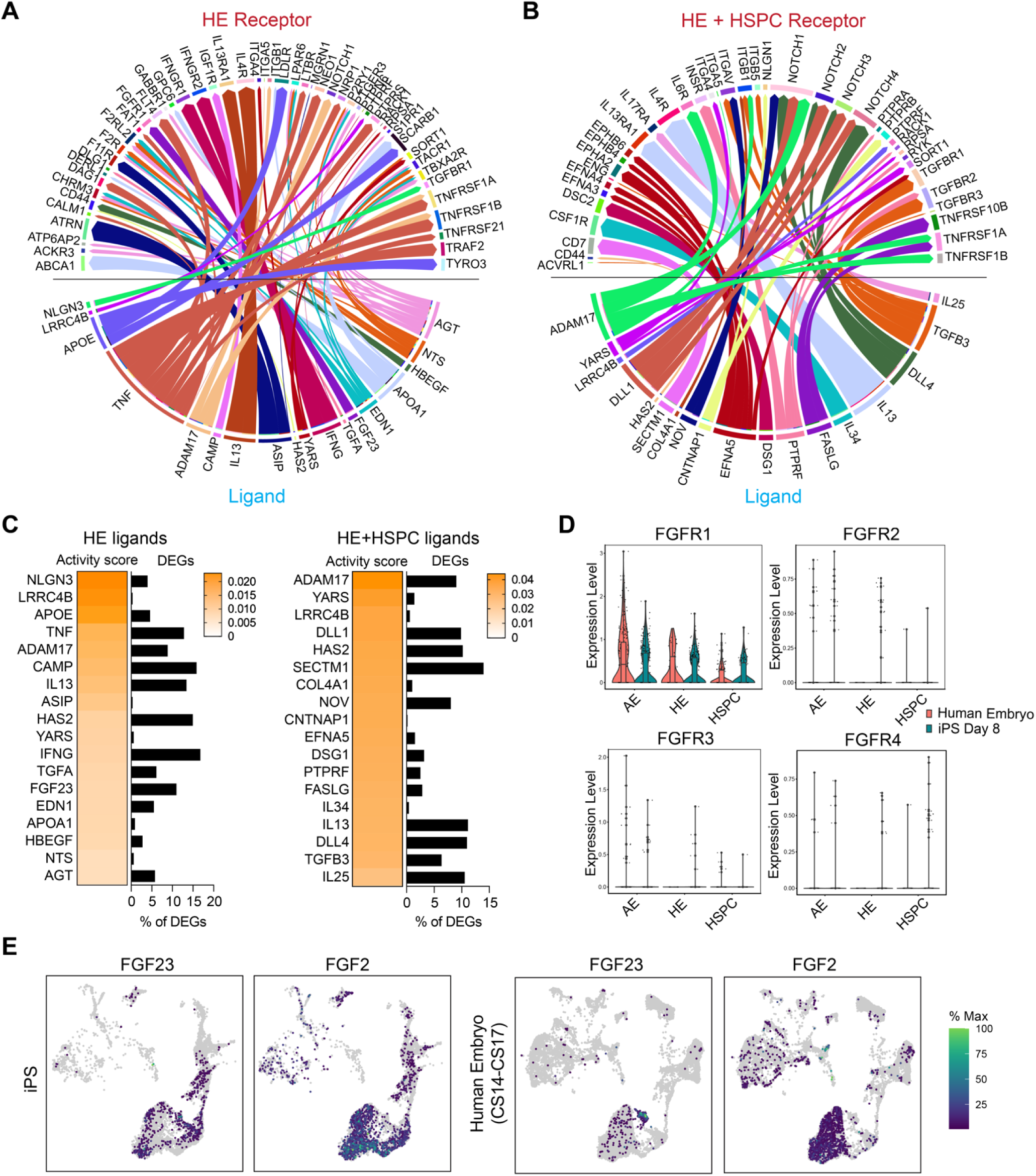
A map of ligand-receptor interactions to inform iPSC differentiation. (**A-B**) Top ligand-receptor interactions identified by NicheNetR v.1 associated with differentially expressed genes in day 8 iPSC EBs vs. *in vivo* (CS14-17 human embryo): (A) hemogenic endothelium (cluster 75 of integrated dataset) and (B) EHT consisting of hemogenic endothelium (cluster 75) and early HSPCs (cluster 74). Ligands (bottom) are connected to their respective receptors (top) expressed on HE or EHT cells, with ligands and receptors separated by a line in the middle. (**C**) Ligands in (A, B) ranked by the activity score defined as the area under the precision recall curve (AUPR) and percent of predicted regulated DEGs. (**D**). Expression of FGF receptors (*FGFR1-4*) during *in vitro* (red) and *in vivo* (blue) EHT. (**E**). Expression of FGF2 and FGF23 ligands in the integrated dataset, split by day 8 iPS (left) or HSC-competent embryo (right).

Endothelial identity was not properly suppressed in iPSC-derived EHT resulting in persistent expression of endothelial genes, such as *KDR* and *FLT1* (**Figure 4I-J, Figure S4J**). Interestingly, fibroblast growth factor FGF23 was identified by NicheNetR analysis as a predicted upstream signal for 36 differentially expressed genes between *in vitro* and HSC-competent *in vivo* HE (**Figure 5A,C, Figure S5A**; **Table S5**). FGF signaling plays an important role in endothelial biology, including by upregulating the expression of *KDR*, and is essential for generation of HSC-competent HE^35–37^. *FGFR1* is the main FGF receptor expressed on HE (**Figure 5D**). Interestingly, iPSC-derived endothelial cells expressed abundant *FGF23* and the more canonical ligand *FGF2* (**Figure 5E**). In addition, *FGF2* was widely expressed in non-endothelial populations in our data (**Figure 5E**). Taken together, these data led us to hypothesize that FGF signaling is attenuated in HSC-competent HE during EHT *in vivo*, but remains activated during iPSC-derived EHT, and that inhibition of FGF signaling during EHT can augment HSPC generation.

### Chemical modulation of FGF signaling during EHT regulates hematopoiesis *in vivo* and *in vitro*

To determine the role of FGF signaling during EHT *in vivo*, we took advantage of the external embryonic development and transparency of zebrafish embryos to monitor FGF activity in real time in the dorsal aorta (DA) using the *dusp6:d2EGFP* transgenic reporter. FGF activity in the DA was attenuated following the onset of EHT at 30 hours post-fertilization (hpf) compared to 21 hpf preceding EHT (**Figure 6A**), suggesting that decreased FGF signaling may be important for EHT. We next temporally modulated FGF signaling by incubating zebrafish embryos with a small-molecule FGFR inhibitor erdafitinib^38^ or an FGF signaling agonist, BCI, that acts by inhibiting a negative feedback loop through DUSP1/6 phosphatases^39^. Erdafitinib and BCI concentrations were titrated using the *dusp6:d2EGFP* reporter to identify concentrations with highest specific activity that did not affect embryonic development (**Figure S6A,B**). Treatment with erdafitinib at 16-24 hpf prior to the onset of EHT did not cause a significant change in the number of *cd41^+^* HSPCs in the floor of the DA at 52 hpf (**Figure S6C,D**). By contrast, treatment with BCI at 16-24 hpf led to a significant decrease in *cd41^+^* HSPCs in the DA, suggesting that FGF signaling is tightly regulated in early HE specification. We next treated zebrafish with erdafitinib and BCI starting at 24 hpf, coinciding with the onset of EHT, and assessed the output of *cd41^+^* HSPCs at 52 hpf. Inhibition of FGF signaling with erdafitinib led to a significant increase in *cd41*^+^ HSPCs in the DA, whereas activation of FGF signaling with BCI led to a significant reduction in *cd41^+^* HSPCs (**Figure 6C,D**). These data demonstrate that FGF signaling is attenuated in HSC-competent HE and negatively regulates EHT, such that chemical inhibition of FGFR during EHT augments HSPC generation *in vivo*.

**Figure 6.**
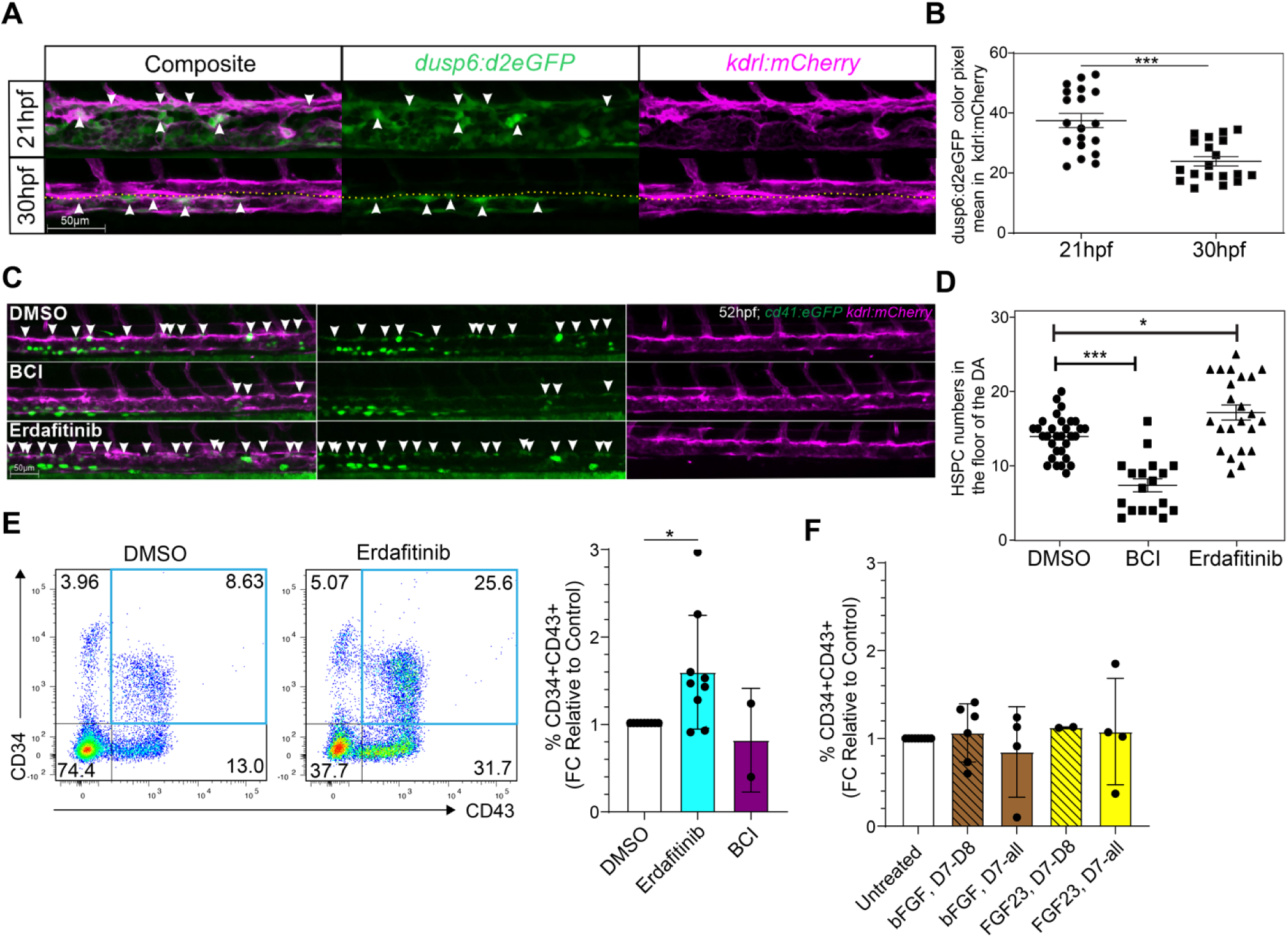
Chemical modulation of FGF signaling during EHT regulates hematopoiesis *in vivo* and *in vitro*. (**A-B**) Representative images (A) and quantification (B) of the number of hemato-endothelial cells with activated FGF signaling (white arrowheads) in zebrafish kdrl+ endothelial cells within the aorta-gonad pronephros prior to EHT (21 hpf) or post-EHT (30 hpf). Yellow dotted line separates the aorta and vein at 30 hpf. At this developmental time, most FGF activity remains in the vein (white arrowheads). Activated FGF signaling is quantitated using the dusp6:d2EGFP; kdrl:mCherry double-positive cells. Mean ± s.e.m. of 30 embryos in 2 independent experiments, unpaired two-tailed Student’s t-test (*p<0.05, ***p<0.001). (**C-D**) Representative confocal images (C) and quantification (D) of the number of HSPCs (white arrows) in the floor of the dorsal aorta (DA) following treatment with an FGF small molecule-inhibitor (erdafitinib) or agonist (BCI). Zebrafish were treated with 7 µM erdafitinib, 10 µM BCI, or vehicle (DMSO) during EHT from 24-52 hpf. *kdrl:mCherry^+^; cd41:eGFP^+^* HSPCs were quantified at 52 hpf. Mean ± s.e.m. of 30 embryos in 2 independent experiments; significance determined by unpaired two-tailed Student’s t-test (*p<0.05, ***p<0.001). (**E**) Representative flow plots (left) and quantification (right) of the number of CD34^+^CD43^+^ iPSC-derived HSPCs following treatment with an FGF small molecule-inhibitor (erdafitinib), agonist (BCI). EBs were treated with 0.1 µM erdafitinib, 10 µM BCI, or vehicle (DMSO) from day 7-8 of differentiation and the proportion of CD34^+^CD43^+^ HSPCs was quantified at day 10. (**F**) The proportion of CD34^+^CD43^+^ iPSC-derived HSPCs following treatment with FGF ligands bFGF and FGF23 or vehicle (0.1% BSA) from days 7-8 (D7-8) or days 7-10 (D7-all) of differentiation. CD34^+^CD43^+^ HSPCs quantified at day 10. (E,F): Mean ± s.d. shown of n = 9 independent experiments with two different iPSC lines for erdafitinib (single outlier point removed), and n = 2 - 6 independent experiments with two different iPSC lines for other conditions; significance determined by paired two-tailed Student’s t-test (*p<0.05).

Lastly, we tested whether chemical inhibition of FGF signaling during iPSC-derived EHT, where FGF activity is predicted to be elevated, can augment HSPC generation from iPSCs. To this end, we treated EBs derived from two different iPSC lines with erdafitinib and BCI starting on day 7 of differentiation, coinciding with the beginning of EHT, and removed the small molecules on day 8. In addition, we tested the addition of bFGF (FGF2) or FGF23 ligands on day 7, with removal on day 8 or continued culture. We then measured the generation of CD34^+^CD43^+^ HSPCs on day 10 of differentiation, 48 hours after the removal of the small molecules to allow for EHT to proceed prior to measuring hematopoietic output. Treatment with 0.1 µM erdafitinib led to a significant increase in the proportion of CD34^+^CD43^+^ HSPCs on day 10 (n = 9 independent experiments) (**Figure 6E**, **Figure S6E**), whereas treatment with FGF agonist BCI, bFGF, or FGF23 ligands had no effect on CD34^+^CD43^+^ HSPC output (**Figure 6F; Figure S6F**). Taken together, we identify FGF signaling as one of the pathways upstream of EHT gene expression and demonstrate that it acts as a negative regulator of EHT both *in vivo* and in the iPSC system. Accordingly, chemical inhibition of FGF signaling during EHT augments iPSC-derived hematopoiesis. Moreover, we identify a map of candidate signaling pathways that can be modulated to improve iPSC differentiation protocols.

## DISCUSSION

EHT is a critical transient stage in hematopoietic ontogeny. Here, we interrogate the single cell kinetics of EHT during iPSC differentiation and integrate it with human embryo datasets to generate a high-resolution map of EHT. Although iPSCs give rise to definitive HE with robust multilineage hematopoietic potential, they do not efficiently generate bona fide HSCs, suggesting broad differences compared to hematopoiesis in the embryo. By comparing the molecular changes at developmentally equivalent stages of EHT *in vitro* and *in vivo*, we uncover several insights. By connecting these expression changes with upstream signaling factors, we identify ligand-receptor pairs predicted to be dysregulated in iPSC differentiation. We identify FGF signaling as one such pathway and demonstrate that it acts as a negative regulator of EHT *in vivo* and in the iPSC system. Chemical inhibition of FGF signaling augments hematopoiesis in the zebrafish and from iPSCs. Our ligand-receptor map provides a resource for future interrogation of pathways that can be modulated to improve iPSC differentiation protocols (**Figure 7**).

**Figure 7.**
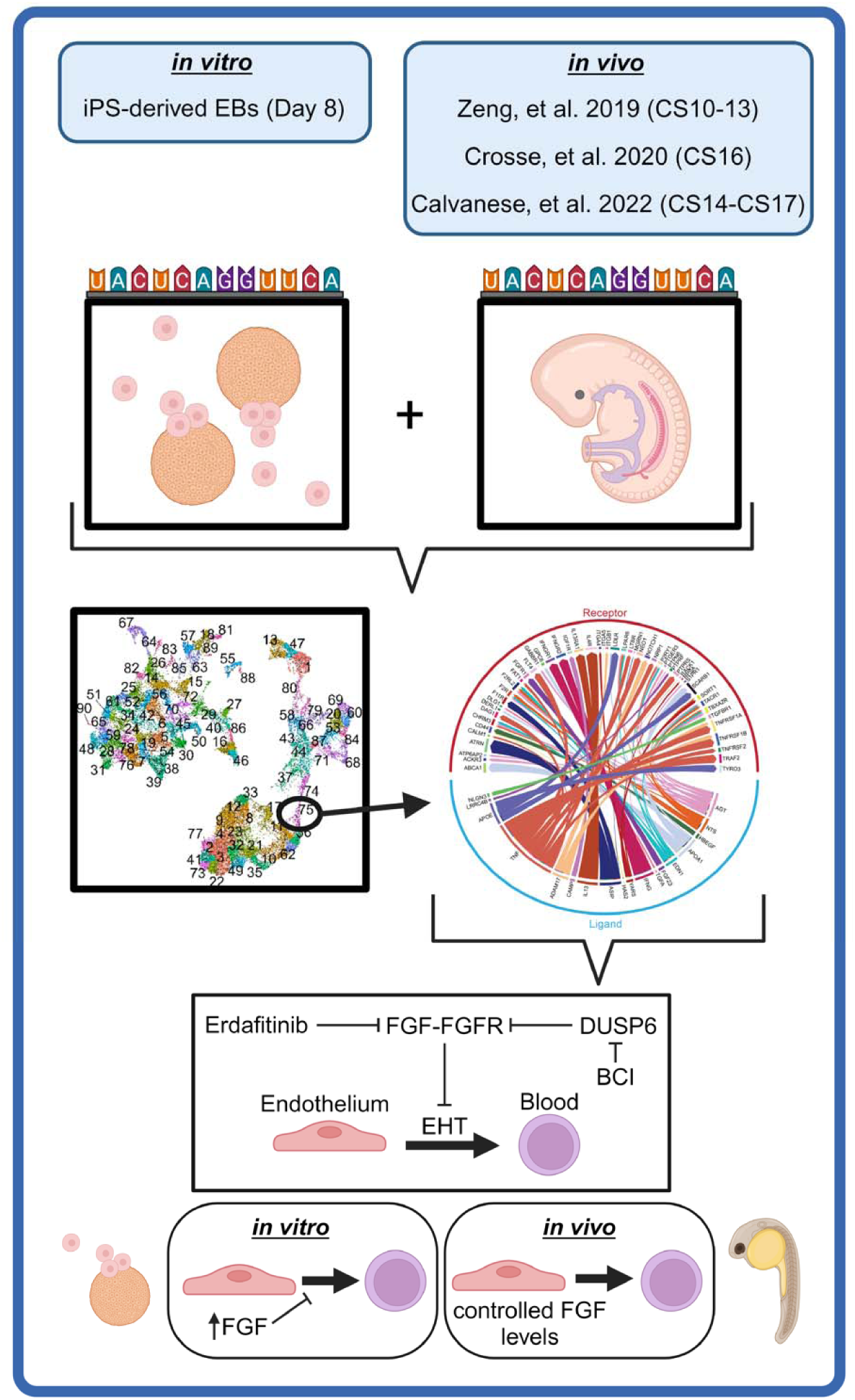
Summary model. We integrate high-resolution single cell profiling of EHT in iPSC-derived with human embryo transcriptomic data to uncover molecular differences and receptor-ligand interactions between *in vitro* and *in vivo* hematopoiesis. The FGF pathway is aberrantly activated during EHT *in vitro* and chemical inhibition of FGF augments zebrafish and iPSC hematopoiesis. Figure created with BioRender.com.

Although several single cell transcriptomic studies have captured iPSC-derived EHT^8,10,15,18,25,26^, we lacked a high-resolution map of gene expression changes during EHT in iPSC differentiation and its direct comparison to the human embryo. To bridge this gap, we first performed an unbiased single cell transcriptional profiling of whole EBs using sci-RNA-seq, which showed that, akin to the embryo, EHT is a transient process that peaks around day 8 of differentiation in our EB system. We then generated a high-resolution map of iPSC-derived EHT at day 8. By comparing the molecular changes at developmentally equivalent stages of EHT *in vitro* and *in vivo*, we uncover several insights. First, as in the embryo, AE clusters closest to HE, suggesting that arterialized endothelium emerges as a precursor to definitive HE in iPSC differentiation, as previously shown^12,18,29,40^. Second, iPSC-derived EHT generates two populations of HE, one of which appears to be a precursor to both erythroid progenitors and a second HE population that acquires an HSC-like signature; however, the expression of key HSC fate genes and signaling factors is diminished overall. Moreover, we find that the developmental transition from AE to HE to HSPC is marked by progressive global silencing of endothelial cell fate and induction of hematopoietic fate. However, we observe a persistent high expression of endothelial fate genes in nascent iPSC-derived HSPCs. Our findings point to the mis-regulation of specific developmental programs in iPSC differentiation. Our integrated dataset is available to the community and can be explored interactively through an online interface (see Code and Data Availability) and will be a resource for further interrogation of EHT dynamics.

EHT in the embryo occurs at specific anatomical sites within developing clusters, critically in the AGM. The nascent HSPCs in these clusters are exposed to defined developmental signals and biophysical forces. By contrast, hematopoiesis in EBs is modulated by signals provided in the culture media. We reasoned that inappropriate signaling pathways in developing EBs can be identified based on expression differences between cells undergoing EHT in iPSCs and human embryos. These may include factors that are present in the embryo and need to be included in the iPSC protocol, or conversely, factors present in iPSC media or secreted in the EBs that are normally absent at sites of hematopoiesis in the embryo. This analysis has uncovered several predicted signaling pathways, for instance TGFα/β, Notch, interferon gamma, and EDN1, known to be essential regulators of EHT^1,34^. We also identified candidate pathways not currently known to regulate EHT or hematopoiesis. Our ligand-receptor analysis provides a resource for future interrogation of pathways to improve iPSC differentiation protocols.

FGF signaling emerged from this analysis as a potential negative regulator of EHT. FGF signaling has previously been identified as essential for specification of HE from endothelial precursors^35,41^. We show that FGF signaling is attenuated at the onset of EHT in the DA, but remains elevated during iPSC-derived EHT. Our analysis has implicated the canonical FGF2 ligand, widely secreted in EBs and commonly supplemented in iPSC differentiation protocols, and a non-canonical FGF23 ligand, linked to adult phosphate and calcium homeostasis with known roles in erythropoiesis, iron regulation and inflammation^42–46^. We show that FGF signaling negatively regulates EHT, consistent with prior reports^47^, and can be chemically modulated to augment hematopoiesis. In the zebrafish, treatment with pan-FGFR inhibitor at the onset of EHT augments HSPC generation, while treatment with a small molecule agonist drastically reduces HSPC production. Similarly, in EB differentiation, inhibition of FGFRs during the developmental window coinciding with the peak of EHT boosts HSPC generation from iPSCs. Given that many iPSC differentiation protocols incorporate exogenous FGFs in the serum or as recombinant proteins^29,48^, the dose and time-dependent modulation of FGF signaling via removal of exogenous FGFs and/or treatment with small molecule inhibitors during the peak of EHT should be considered to optimize HSPC generation in iPSC differentiation protocols, and perhaps optimize rare HSC fate specification *in vitro*.

In summary, single cell comparative analysis of EHT and ligand-receptor predictions identify a negative role of FGF signaling in hematopoietic ontogeny. Our single cell map of EHT and ligand-receptor analysis provide a resource for future interrogation of novel pathways to improve iPSC differentiation protocols.

## METHODS

### Human iPSC lines

Normal iPSC lines MSC-iPSC1^49^ and CD45-iPSC1^50^ were maintained on hESC-Qualified Matrigel (Fisher Scientific, #8774552) in mTESR Plus media (Stem Cell Technologies, #100-0276). Prior to EB formation, iPSCs were transferred onto mouse embryonic fibroblasts (MEFs) and cultured in iPSC-MEF media: DMEM/F12 (Thermo Fisher, #11330-032) supplemented with 20% knockout serum replacement (KOSR; Thermo Fisher, #10828028), 1 mM L-glutamine (Thermo Fisher, #25030081), 1X non-essential amino acids (MEM-NEAA; Thermo Fisher, #11140-050), 55 µM 2-mercaptoethanol (Thermo Fisher, #21985-023), and 10 ng/mL basic fibroblast growth factor (bFGF, Peprotech #100-18C). MEFs were cultured in MEF media consisting of DMEM (Thermo Fisher, #11960069), 9.1% batch tested fetal bovine serum (Sigma #F0926, lot: 201M111), 1 mM L-glutamine, 1X MEM-NEAA, and 1 mM sodium pyruvate (Thermo Fisher, cat# 111360-070) and treated with mitomycin C (Thermo Fisher #BP25312) prior to use.

### Embryoid body differentiation

MEF-cultured iPSCs were treated with 1 mg/mL collagenase IV (Thermo Fisher, cat# 17104019) prior to scraping whole colonies. Scraped whole colonies were allowed to gravity settle then were transferred into EB media: KO DMEM (Thermo Fisher, #10829-018), 20% FBS, 1 mM L-glutamine, 1X MEM-NEAA, 55 µM 2-mercaptoethanol, 50 μg/mL ascorbic acid (Sigma, #A4544-25G), 100 U/mL penicillin streptomycin (Thermo Fisher, #15140122), and 100 μg/mL holo-transferrin (Fisher Scientific, #61-639). Colonies were put on a shaker at 37°C (Day 0 of differentiation, a.k.a. D0). After approximately 24hr (D1), EBs were collected into 15 mL conical tubes and gravity settled. They were resuspended in EB culture media supplemented with final concentrations of 100 ng/mL human stem cell factor (hSCF, Peprotech #300-07), 100 ng/mL fms-like tyrosine kinase 3 ligand (hFLT3L, Peprotech #300-19), 50 ng/mL bone morphogenic protein 4 (hBMP4, R&D Systems #314-BP), 10 ng/mL granulocyte colony-stimulating factor (hG-CSF, Peprotech #300-23), 25 ng/mL interleukin-6 (hIL-6, Peprotech #200-06) and 10 ng/mL interleukin-3 (hIL-3, Peprotech #200-03) and again put on a shaker at 37°C. EB culture media with the listed cytokines was refreshed on D5 and D9. On days of collection for sc-RNA-seq or flow cytometry, EBs and all surrounding media were collected into 15 mL conical tubes and spun at 1350 rpm for 5 minutes to ensure the capture of both cells within the EBs as well as floating hematopoietic cells. The samples were then incubated in IMDM (Thermo Fisher, #12440061) with 2 mg/mL Collagenase B (Sigma-Aldrich, #11088807001) at 37°C for 2 hours while shaking with trituration approximately every 30 minutes. After 2 hours, 9 mL of IMDM was added to the EB solution and the samples were spun at 1350 rpm for 5 minutes. The EBs were then incubated in 2 mL Gentle Cell dissociation reagent (Stem Cell Technologies, #100-0485) in a 37°C water bath for 15 minutes followed by vigorous trituration until they formed a single cell solution. 10 mL of 3% fetal bovine serum (FBS) in PBS was added and the solution was filtered through a 70 µm filter with a subsequent 2 mL 3% FBS in PBS wash. The single cell solution was then spun down at 1350 rpm for 5 minutes, resuspended in 3% FBS in PBS and the cells were counted using a Denovix CellDrop cell counter prior to subsequent assays.

### Zebrafish husbandry and strains

All experiments in zebrafish (*Danio rerio*) were performed according to the Guidelines for Ethical Conduct in the Care and Use of Animals^51^, Iowa State University Animal Care and Use Committee IACUC-20-025, and IACUC-20-024 approved protocols, and in compliance with ARRIVE guidelines^52^, and the American Veterinary Medical Association (2020) and NIH guidelines for the humane use of animals in research.

*Tg*(*dusp6:d2EGFP*)^pt6^ zebrafish were kindly donated by Michael Tsang^53^. Other zebrafish lines used in this study were: wt AB (ZIRC), *Tg(kdrl:HsHRAS-mCherry)s896* ^4^ (referred to as *kdrl:mCherry* throughout the manuscript), *Tg(−6.0itga2b:eGFP)la2*^54^ (referred to as *cd41:eGFP* throughout manuscript), and various intercrosses of these lines were utilized. Zebrafish were mated, staged, raised, and processed as described^55^ in a circulating aquarium system at 28°C.

### Zebrafish chemical treatments and microscopic visualization of *Tg(dusp6:d2EGFP)*

30 16 hpf heterozygous *Tg(dusp6:d2EGFP)* zebrafish embryos were placed per well on 12-well plates containing 3 mL embryo water plus 5 μM or 7 μM Erdafitinib (JNJ-42756493, Item No. 21813, Cayman Chemical); or 5 μM or 20 μM BCI (hydrochloride) (Item No. 21945, Cayman Chemical). DMSO was added along with the chemicals to reach a final concentration of 1% to increase permeability. Chemicals were washed off with embryo water at 24 hpf. Embryos were then dechorionated with pronase (11459643001, Sigma), anesthetized in 200 mg/ml tricaine (Syncaine® Fish Anesthetics, MS-222) and the anterior part of the embryo was imaged for dusp6 activity in a Leica M205 FCA with Leica Application suite X software (v3.7.6.25997).

### Live confocal imaging of HSPCs and *dusp6* activity in zebrafish embryos

To visualize HSPCs and *dusp6* activity, live confocal microscopy was performed at 52 hpf on *Tg(cd41:eGFP*; *kdrl:mCherry),* and *Tg(dusp6:d2EGFP; kdrl:mCherry)*, double-transgenic embryos, respectively, at the indicated developmental times. Z sections of the DA region were imaged on a Zeiss LSM 700 Laser Scanning Confocal with Zen Black software (v14.0.27.201). *dusp6* activity within the *kdrl:mCherry^+^* cells of the aorta-gonad-pronephros region of the embryo was quantified from individual Z stacks by ImageJ (v1.53f51), by enclosing 3-4 *kdrl^+^* endothelial ROI’s limited by two contiguous intersegmental vessels using the “Freehand selection”. Green channel pixel mean values were then analyzed using “Color Histogram”. *cd41^+^, kdrl^+^* HSPCs were manually counted throughout the confocal stack comprising the DA and denoted by arrows. Images were processed with ImageJ (v1.53f51). All animals were anesthetized in 200 mg/ml tricaine (Syncaine® Fish Anesthetics, MS-222) before microscopic analysis.

### Flow cytometry

For the day 8 10X sc-RNA-seq experiment, EBs derived from the 45-iPSC line were dissociated as described above and stained with the following antibodies (all BD Biosciences): CD34-PE (#555822), CD43-PerCpCy5.5 (#563521), CD45-APC (#555485). DAPI (ThermoFisher, #D1306) was used as a live-dead stain for all experiments. The cells were sorted on a BD FACSAria III into 50% FBS in PBS. For HSPC quantification, 100,000 EB dissociated cells were stained with the following antibodies (all BD Biosciences): CD34-PeCy7 (#560710), CD144-FITC (#560411), CD43-PerCpCy5.5, CD45-APC, and DLL4-PE (#564412) in 3% FBS in PBS. Samples collected on days 5-8 were stained as above with CD184-APC (#560936) replacing CD45-APC. A single sample was used for isotype staining.

### Single cell RNA sequencing

For single cell combinatorial indexing RNA-sequencing (sci-RNA-seq), EBs derived from both MSC-iPSC and 45-iPSC were dissociated on days 7, 8, 11, 14, 18, and 21 per the protocol outlined above. Cells were then resuspended in a cell lysis buffer (10mM Tris-HCl pH 7.4, 10mM NaCl, 3mM MgCl2, 0.1% IGEPAL CA-630) followed by gentle pipetting. The resulting nuclei were then filtered through a 70 μm strainer and incubated in 4% paraformaldehyde in 1.25X PBS for 15 minutes with inversion every 5 minutes. The nuclei were then spun at 800g for 6 minutes and resuspended in nuclei suspension buffer (10mM Tris-HCl pH 7.4, 10mM NaCl, 3mM MgCl2, 1% SUPERase In RNase Inhibitor). The nuclei were flash frozen and stored in liquid nitrogen until processing. These nuclei samples were submitted to the Brotman Baty Institute (BBI) for sci-RNA-seq (3-level, RNA3-028-a), which was performed similarly to the published protocol (http://dx.doi.org/10.17504/protocols.io.9yih7ue) and sequenced using a NextSeq 2000. For the 10X 3’ Chromium Next GEM v. 3.1 sc-RNA-seq sample, 45-iPSC-derived day 8 EBs were dissociated and FACS sorted according to the protocol outlined above and were spun down and resuspended in 0.04% ultrapure BSA post-sort, then loaded onto a Chromium single cell controller. All subsequent steps were performed according to the manufacturer’s protocol. Sequencing was performed on a NovaSeq 6000 S1.

### FASTQ file processing

FASTQ files for the sci-RNA-seq samples were used to generate barcode, feature, and count matrix files internally by the BBI according to their processing pipeline (see published protocol above). FASTQ files for the 10X iPSC-derived EB sample were processed using CellRanger v.4.0.0 to align reads to the GRCh38-2020-A reference genome. FASTQs from Zeng et al.^8^ were also processed using CellRanger v.4.0.0 to align reads to the GRCh38-2020-A reference genome. FASTQs from Crosse et al.^9^ were processed using CellRanger v.3.1.0 on the 10X Cloud to align to the GRCh38-3.0.0 reference genome. The count matrices and sample information submitted to the GEO database by Calvanese et al.^10^ were used for downstream analyses.

### Sci-RNA-Seq transcriptomic analysis

The Monocle 3 v1.3.1 R package^56^ was used for the analysis of sci-RNA-seq data. Cells with a low number of unique molecular identifiers (UMIs) were removed (see GitHub, linked in the Code and Data Availability section, for the code and cutoffs used). The maximum number of UMIs detected across all samples was 62698, so no upper UMI limit was set. Cells with more than 10% of UMIs mapping to mitochondrial genes were also removed. All samples were combined into a single cell_data_set object for downstream analysis. Batch correction was not needed.

### 10X 3’ sc-RNA-seq transcriptomic analysis

The Seurat v.4.0.1 R package^57^ was used for the initial analysis of 10X sc-RNA-seq data. Density plots were utilized to determine all UMI and percent mitochondrial read cutoffs. Cells with UMIs < 100, UMIs > 100,000 and more than 20% mitochondrial reads were removed from the day 8 iPSC-derived EB dataset. Cells with UMIs < 100, UMIs > 7000, and more than 20% mitochondrial reads were removed from the Zeng et al. CS10 data. Cells with UMIs < 100, UMIs > 9000, and more than 10% mitochondrial reads were removed from the Zeng et al. CS11 data. Cells with UMIs < 100, UMIs > 7000, and more than 40% mitochondrial reads were removed from the Zeng et al. CS13 data. Cells with UMIs < 100, UMIs > 100,000, and more than 10% mitochondrial reads were removed from all Crosse et al. data. Cells with UMIs < 500 and more than 5% mitochondrial reads were removed from the Calvanese et al. data. Each dataset was normalized individually prior to integration. As the canonical correlation analysis (CCA) integration function in Seurat requires a selection of top variable features, the FindVariableFeatures function was set to the maximum number of features in each dataset to avoid filtering of relevant genes as setting the maximum number any lower than the maximum number of genes represented in the dataset resulted in improper alignment (data not shown). The datasets were then subsequently integrated using CCA. The integrated data was dimensionally reduced using 50 principal components and two-dimensional unique manifold approximation projection (UMAP). For downstream differential gene expression and pseudotime analysis, the Seurat integration object was converted into a Monocle cell_data_set object for use in Monocle 3 pipelines.

### Differential expression analysis

All differential expression analysis (except that conducted internally by NicheNetR) was conducted using Monocle 3^56^. For the comparison of *in vitro* to *in vivo* HE or EHT populations, a mixed-negative binomial model to remove batch effects associated with dataset and/or stage (categorized as iPS, Crosse et al. or Calvanese et al., or CS10-17 or iPS, respectively) was used. To identify genes that were differentially expressed between AE (Monocle 3 cluster 36) and HE (Monocle 3 cluster 75), or HE (Monocle 3 cluster 75) and early HSPCs (Monocle 3 Cluster 74) for the day 8 iPSC-derived EB *in vitro* data, a standard linear regression as implemented by Monocle 3 defaults was utilized since there were no batch effects associated with dataset to be removed. To identify genes that were differentially expressed between AE (Monocle 3 cluster 36) and HE (Monocle 3 cluster 75), or HE (Monocle 3 cluster 75) and early HSPCs (Monocle 3 Cluster 74) for the HSC-competent *in vivo* data (Crosse et al. and Calvanese et al.), a mixed-negative binomial model to remove batch effects associated with dataset (publication) was used. Differentially expressed genes with q-value > 0.05 and -0.25 < estimates (log_2_ Fold Change) < 0.25 were filtered out.

### Cell type identification

Hematopoietic-related populations in the sci-RNA-seq data were identified by generating aggregate expression scores for sets of genes identified by Xu et al. for each cell type, which were then compiled into relative expression (scaled) heatmaps. These scores were used to annotate Monocle 3-determined clusters as specific cell types based on the maximal expression score for each cluster. See the provided GitHub repository in the Code and Data Availability section for code pertaining to the generation of the aggregate gene set scores and cluster resolution. Similarly, day 8 iPSC 10X cell types were annotated accordingly with the aggregate gene expression score of cell type gene sets from either Zeng et al. or Calvanese et al.

### Gene Ontology

The list of differentially expressed genes (q < 0.05, 0.25 < Log_2_(FC) < -0.25) from the comparison of *in vitro* to *in vivo* HE or EHT populations was uploaded to Metascape v.3.5^58^ for Gene Ontology (GO) analysis. Significant (q-value / FDR < 0.05) summary GO Biological Process Terms were filtered from the resulting output files. To compare pathways that are differentially regulated during the AE-HE or HE-HSPC transitions between *in vitro* and HSC-dependent *in vivo*, the differentially expressed gene lists for *in vitro* and *in vivo* were uploaded together to Metascape for batch analysis. Significant (q-value / FDR < 0.05) summary GO biological process terms were then identified.

### NicheNetR analysis

For NicheNetR analysis, NicheNetR^33^ (12-01-2022 released version) functions were used that internally implemented the Wilcoxon-rank sum for differential expression analysis via Seurat v.4, which identified a similar, although more limited, set of differentially expressed genes (compared to the Monocle 3 linear regression analyses) that was ultimately used for upstream ligand-receptor identification (**Table S5**). Analysis was done following the NicheNetR Seurat vignette. Receiver cells were designated as either HE (Seurat cluster 61) or EHT populations (Seurat cluster 61 and Seurat cluster 59). Sender cells were set as undetermined to reflect the unknown nature of the source of the ligands that may be signaling to the HE or EHT populations in our data. See the GitHub repository linked in the Code and Data Availability section for the complete code.

### FGF treatment of iPSC-derived EBs

For erdafitinib, BCI and FGF ligand treatments *in vitro*, EBs were generated following the same protocol outlined above. On day 7, EBs were pooled in a 50 mL conical tube and split as evenly as possible into each of the different conditions. Erdafitinib (MedChemExpress, #HY-18708) was titrated and used at 0.1 µM as described previously^38^. 10 µM BCI (Cayman Chemicals, #95130-23-7) was used as in the zebrafish experiments. Both erdafitinib and BCI were removed approximately 24 hours after addition. For treatment with FGF ligands: 10 ng/mL bFGF (Peprotech, #100-18C) or 100 ng/mL FGF23 (Peprotech, #100-52) was added. Erdafitinib and BCI were both stored at 1000X concentration in DMSO (Santa Cruz Biotechnology, #67-68-5). bFGF and FGF23 were stored at 1000X concentration in 0.1% BSA in PBS. For the DMSO control, 1 µL of DMSO was added per mL of media. For the untreated control, 1 µL of 0.1% BSA (Thermo Fisher, #15260-037) in PBS was added per mL of media. On day 10, EBs were dissociated as described above and the percentage of live CD34^+^CD43^+^ HSPCs was defined by flow cytometry. The percentage of CD34^+^CD43^+^ HSPCs in the erdafitinib and BCI treatments were normalized to the DMSO vehicle control for each experiment. The percentage of CD34^+^CD43^+^ HSPCs in the bFGF or FGF23 treatments were normalized to the untreated condition. If multiple wells were collected for each condition in an experiment (technical replicates), the average percentage was calculated for each condition prior to normalization.

### Statistical analysis

For iPSC experiments, statistical analysis was performed using GraphPad Prism 10. Paired, two-tailed Students t-test was used for normalized control and treatment groups with a significance cut-off of p < 0.05. For zebrafish experiments, GraphPad Prism 5 was utilized to perform statistical analysis and represent data. Data were analyzed by unpaired two-tailed t-test and confidence intervals at 95%, or ordinary one-way ANOVA with Tukey’s post-test and confidence intervals at 95%.

## Code and Data Availability

All code used to generate figures and data for this manuscript can be found online at https://github.com/FredHutch/Wellington-et-al-2024. The iPSC-derived and integrated single cell datasets can be explored interactively through the CZ CELLxGENE Discover portal: https://cellxgene.cziscience.com/collections/4a2c25af-558a-45fc-bc9a-54ec44a1d63f. All sequencing files generated as part of this study can be found in the Gene Expression Omnibus under accessions GSE274082 (sci-RNA-seq) and GSE274084 (10X sc-RNA-seq). Sequencing data for Zeng et al.^8^ was downloaded from GSE135202. Sequencing data for Crosse et al.^9^ was downloaded from GSE151877. Sequencing data for Calvanese et al.^10^ was downloaded from GSE162950.

## Acknowledgements

We thank the Brotman Baty Institute, particularly Cailyn Spurrell, for generating the sci-RNA-seq data. We thank the Chan-Zuckerberg Initiative for hosting our data on their CZ CELLxGENE Discover portal (https://cellxgene.cziscience.com). We thank Takashi Ishida, Adam Heck, Eliza Barkan, Aishwarya Gogate, Sanjay Srivatsan, David Read, Jennifer Franks and Amy Tresenrider for help with elements of the code used during analysis. We thank Katie Kraskouskas, Stacey Dozono and Sintra Stewart for their assistance in iPSC culture maintenance. We thank Janis Abkowitz, Hannele Ruohola-Baker and Lea Starita for advice throughout the generation of the data in this manuscript. Aurelio Silverstroni (Pathology Flow Core) for technical assistance. S.D. is supported by the NIH/NHLBI (R01 HL151651), NIH/NHLBI (R01 HL169156), NIH/NIDDK (RC2 DK127989), NIH New Innovator Award (DP2 HL147126), Wayne D. Kuni and Joan E. Kuni Foundation Discovery Grant, Edward P. Evans Foundation Discovery Research Grant. S.D. is a Scholar of the Leukemia and Lymphoma Society (1391-24). B.H. is supported by the NIH/NHLBI (R01 HL168110). R.W. is supported by the Institute for Stem Cell and Regenerative Medicine (ISCRM) fellows award. R.E-P is supported by the NIH-NIDDK R01 (R01DK131162), and the Roy J. Carver Charitable Trust (#21–5532).

## Author Contributions

R.W. performed all single cell analysis and iPSC experiments. X.C. performed all zebrafish experiments. R.W., X.C., S.D., B.H., R.E-P., C.T., and C.C., analyzed data and/or assisted with data interpretation. R.W., B.H. and S.D. wrote the paper.

## Competing Interests

CT is a scientific advisory board member, consultant, and/or cofounder of Algen Biotechnologies, Altius Therapeutics, and Scale Biosciences. The other authors declare no competing interests.

## SUPPLEMENTAL TABLE LEGENDS

**Table S1. S1A,B**: Aggregate gene scores for each cluster in Figure 1 and Figure S1D based on embryonic cell type markers from Xu et al.^27^ (Table S1A-B). **S1C**: Hematopoietic and endothelial scores for each cell in the dataset based on the expression of hematopoietic (*RUNX1*, *SPN*, *PTPRC*) or endothelial (*CDH5*, *KDR*, *TIE1*, *CD34, hematopoietic aggregate score = 0*) lineage specific genes. **S1D**: Marker genes from Xu et al.^27^ used to calculate aggregate gene scores in S1A and S1B. Metadata and unprocessed data for the sci-RNA-seq dataset is available from GEO under accession GSE274082.

**Table S2**. Metadata corresponding to the 10X sc-RNA-seq dataset of day 8 iPSC: barcodes, UMI numbers, UMAP coordinates, cluster assignments, Zeng et al.^8^ cell type scores and Calvanese et al.^10^ cell type scores for all cells post-filtering. The markers used for calculating the aggregate gene scores for cell types from Zeng et al. and Calvanese et al. are included in S2B. Related to Figure 2 and Figure S2. Metadata and unprocessed data for the 10X dataset is available from GEO under accession GSE274084.

**Table S3.** Metadata corresponding to the 10X day 8 iPSC, Zeng et al.^8^, Crosse et al.^9^, and Calvanese et al.^10^, integrated dataset: barcodes, dataset, UMI numbers, UMAP coordinates, cluster assignments, and transferred labels from Zeng et al., Crosse et al., and Calvanese et al. for all cells post-filtering in the integrated sc-RNA-seq dataset from Figure 3.

**Table S4A.** Differentially expressed genes (DEGs) for the following comparisons. **S4A-HE**: iPSC day 8 v. CS14-CS16 HE (cluster 75) (HE was not present during CS17). **S4A-EHT**: iPSC day 8 v. CS14-CS17 EHT (clusters 75 and 74). **S4A-HSPC**: iPSC day 8 v. CS14-CS17 early HSPCs (cluster 74). **S4A-invivo-AE-HE**: CS14-CS17 AE (cluster 36) v. CS14-CS16 HE (cluster 75). **S4A-invitro-AE-HE**: iPSC AE (cluster 36) v. iPSC HE (cluster 75). **S4A-invivo-HE-HSPC**: CS14-CS16 HE (cluster 75) v. CS14-CS17 early HSPCs (cluster 74). **S4A-invitro-HE-HSPC**: iPSC HE (cluster 75) v. iPSC early HSPCs (cluster 74). Related to Figure 4 and Figure S4. For all comparisons, q < 0.05, -0.25 > log_2_ fold change > 0.25.

**Table S4B.** Metascape GO term enrichment statistics and associated genes for the comparisons in Table S4A. **S4B-HE**: iPSC day 8 v. CS14-CS16 HE (cluster 75) (HE was not present during CS17). **S4B-EHT**: iPSC day 8 v. CS14-CS17 EHT (clusters 75 and 74). **S4B-HSPC**: iPSC day 8 v. CS14-CS17 early HSPCs (cluster 74). **S4B-AE-HE**: Metascape comparisons in batch mode of CS14-CS17 AE (cluster 36) v. CS14-CS16 HE (cluster 75) and iPSC AE (cluster 36) v. iPSC HE (cluster 75) DEG-enriched GO terms. GO terms for upregulated and downregulated genes shown in separate tabs. **S4B-HE-HSPC**: Metascape comparisons in batch model of CS14-CS16 HE (cluster 75) v. CS14-CS17 early HSPCs (cluster 74) and iPSC HE (cluster 75) v. iPSC early HSPCs (cluster 74) DEG-enriched GO terms. GO terms for upregulated and downregulated genes shown in separate tabs. Related to Figure 4 and Figure S4.

**Table S5.** NicheNetR-identified ligand-receptor pairs associated with differentially expressed genes in HE (cluster 75, iPSC day 8 vs CS14-CS16) and EHT (cluster 75+74, iPSC day 8 vs CS14-CS17). Related to Figure 5 and Figure S5. **S5A**: Ligand-receptor pairs and prior interaction potential for HE. **S5B**: Ligand-receptor pairs and prior interaction potential for EHT. **S5C**: Regulatory scores of upstream ligand-receptors for individual DEGs as calculated by NicheNetR. Related to Figure S5A (HE). **S5D**: Regulatory scores of upstream ligand-receptors for individual DEGs as calculated by NicheNetR. Related to Figure S5B (EHT).

## SUPPLEMENTAL FIGURES

**Figure S1.**
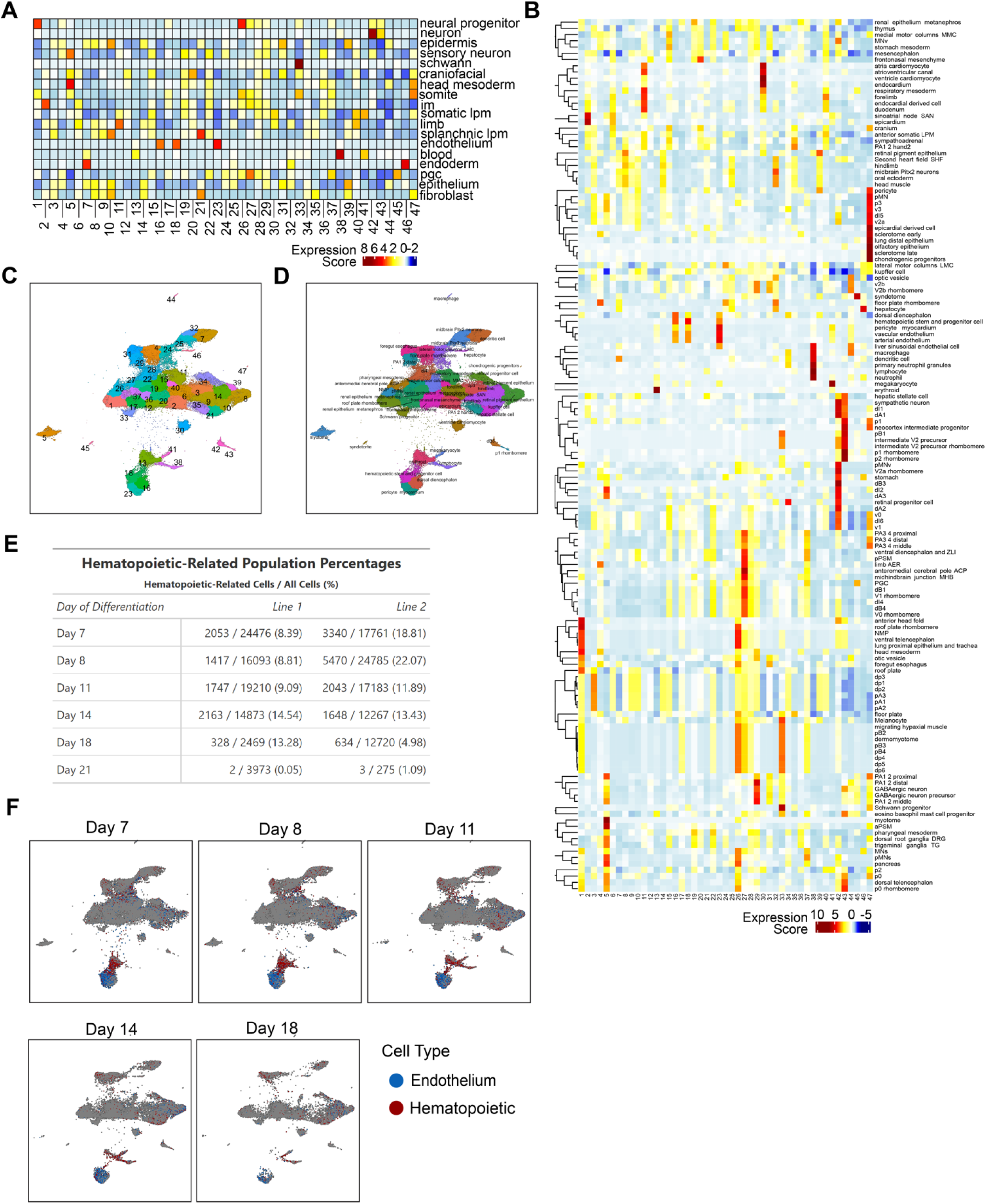
Expanded single cell temporal profiling of hematopoietic ontogeny *in vitro*. Related to Figure 1. (**A**) Cell type scoring of sci-RNA-seq clusters based on Xu et al.^27^ cell type markers (Table S1). (**B**) Cell type scoring of sci-RNA-seq clusters based on extended Xu et al.^27^ cell type markers (Table S1). (**C**) Unsupervised clustering of the sci-RNA-seq dataset using Monocle 3. (**D**) sci-RNA-seq clusters annotated with the maximum cell type score from (B). (**E**) Proportion of hematopoietic cells in the sci-RNA-seq dataset by day of differentiation; two different normal iPSC lines. (**F**) Hematopoietic (red) and endothelial (blue) populations shown by day of differentiation showing the temporal dynamics of hematopoiesis. Cell types were identified as hematopoietic (*RUNX1*, *SPN*, *PTPRC* aggregate gene score) or endothelial (*CDH5*, *KDR*, *TIE1*, *CD34*, and zero hematopoietic aggregate gene score).

**Figure S2.**
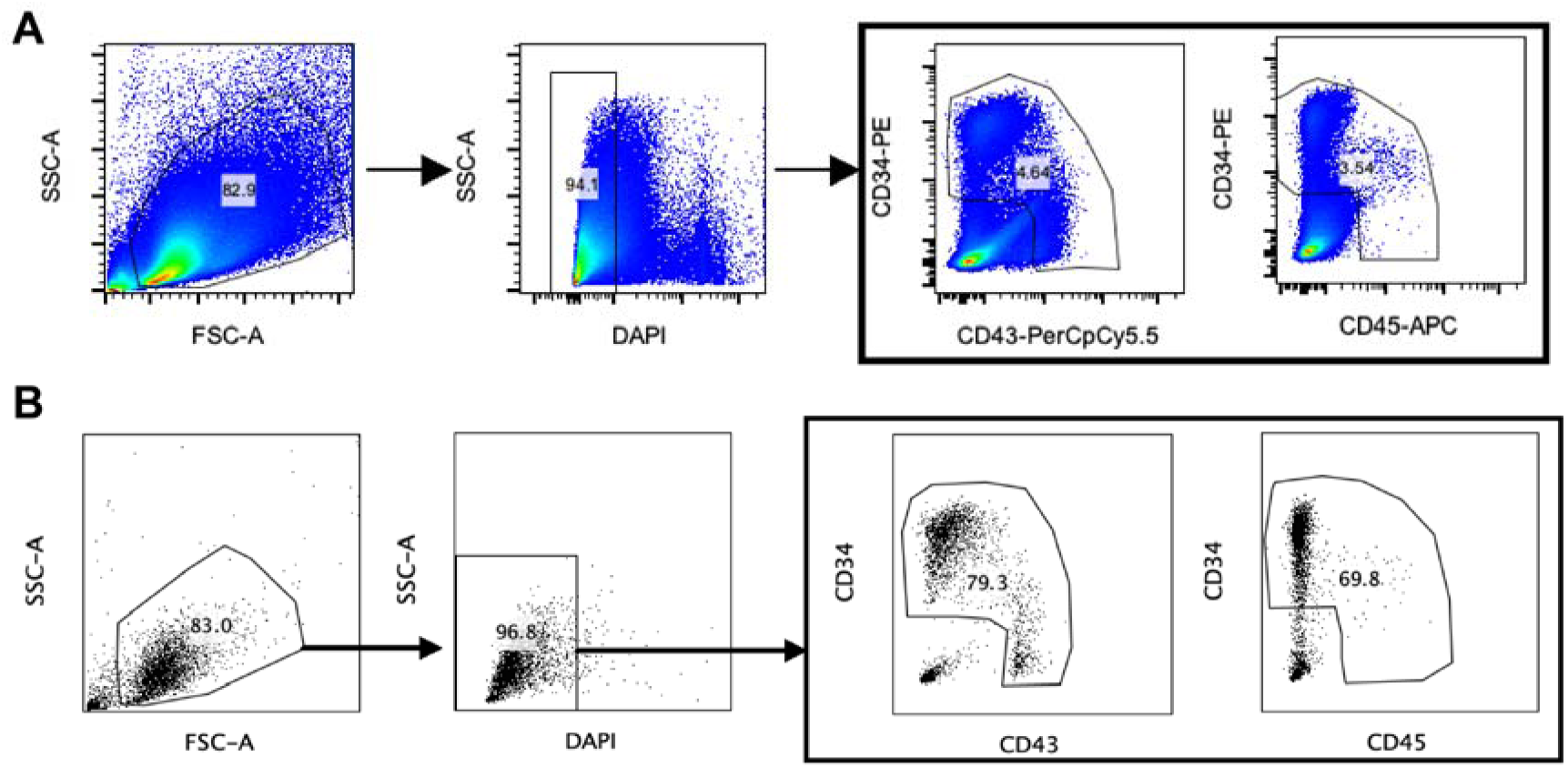
Isolation of iPSC-derived hematopoietic and endothelial cells for 10X sequencing. Related to Figure 2. (**A**). Flow sort gating for isolation of CD34^+^CD43^+/-^CD45^+/-^ hematopoietic and endothelial cells from day 8 of EB differentiation. (**B**) Flow sorted populations from (A) showing sorting purity of hematopoietic and endothelial populations.

**Figure S3.**
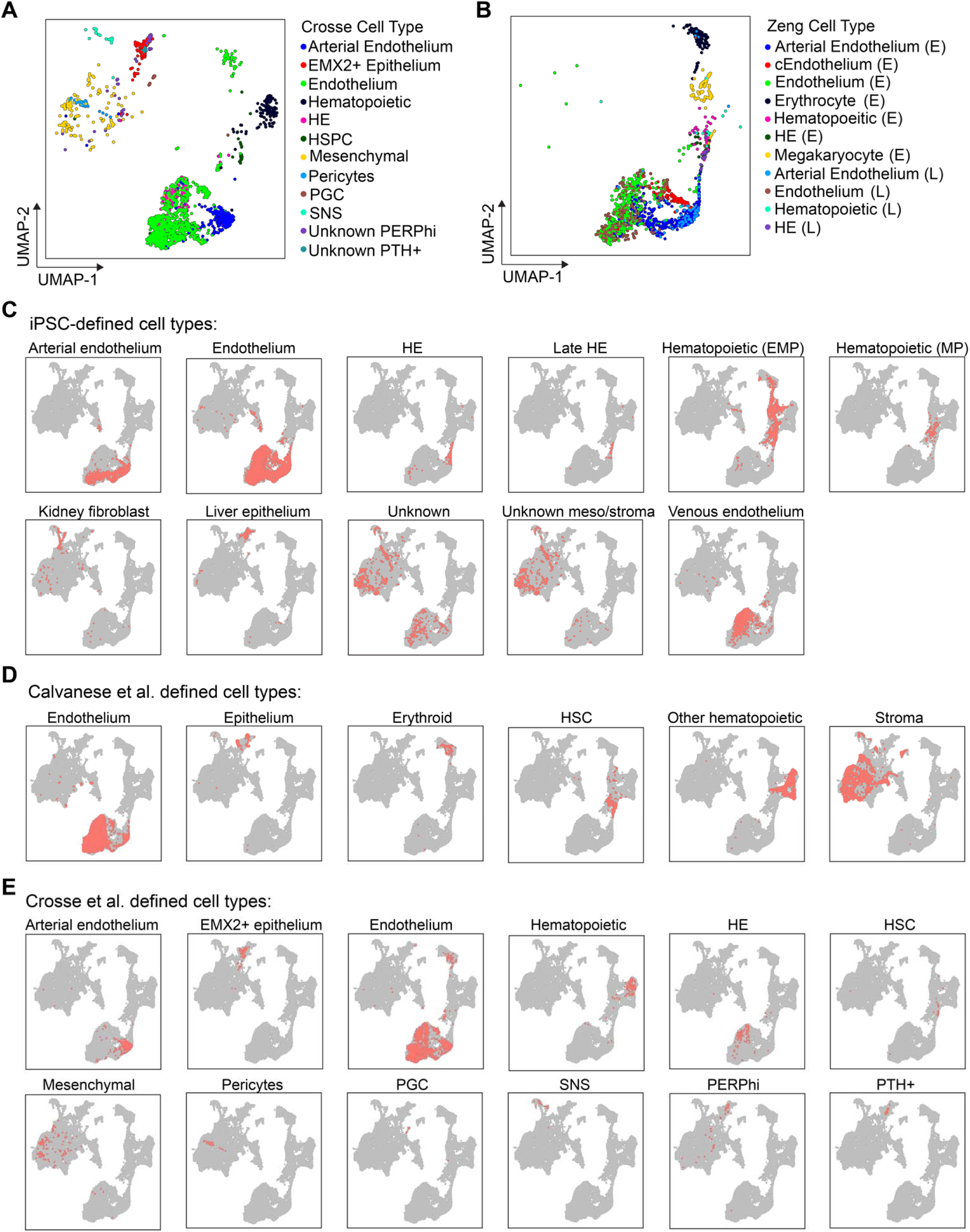
Cell type label transfers for validation of the integrated dataset. Related to Figure 3. (**A**) Cell type label transfer from Crosse et al.^9^ onto the integrated scRNA-seq dataset. (**B**) Cell type label transfer from Zeng et al.^8^ onto the integrated scRNA-seq dataset. (**C-E**) Label transfer of key developmental populations from the original day 8 iPSC (C), Calvanese et al.^10^ (D), and Crosse et al.^9^ (E) datasets onto the integrated UMAP to determine where previously identified cell types are in integrated UMAP space.

**Figure S4.**
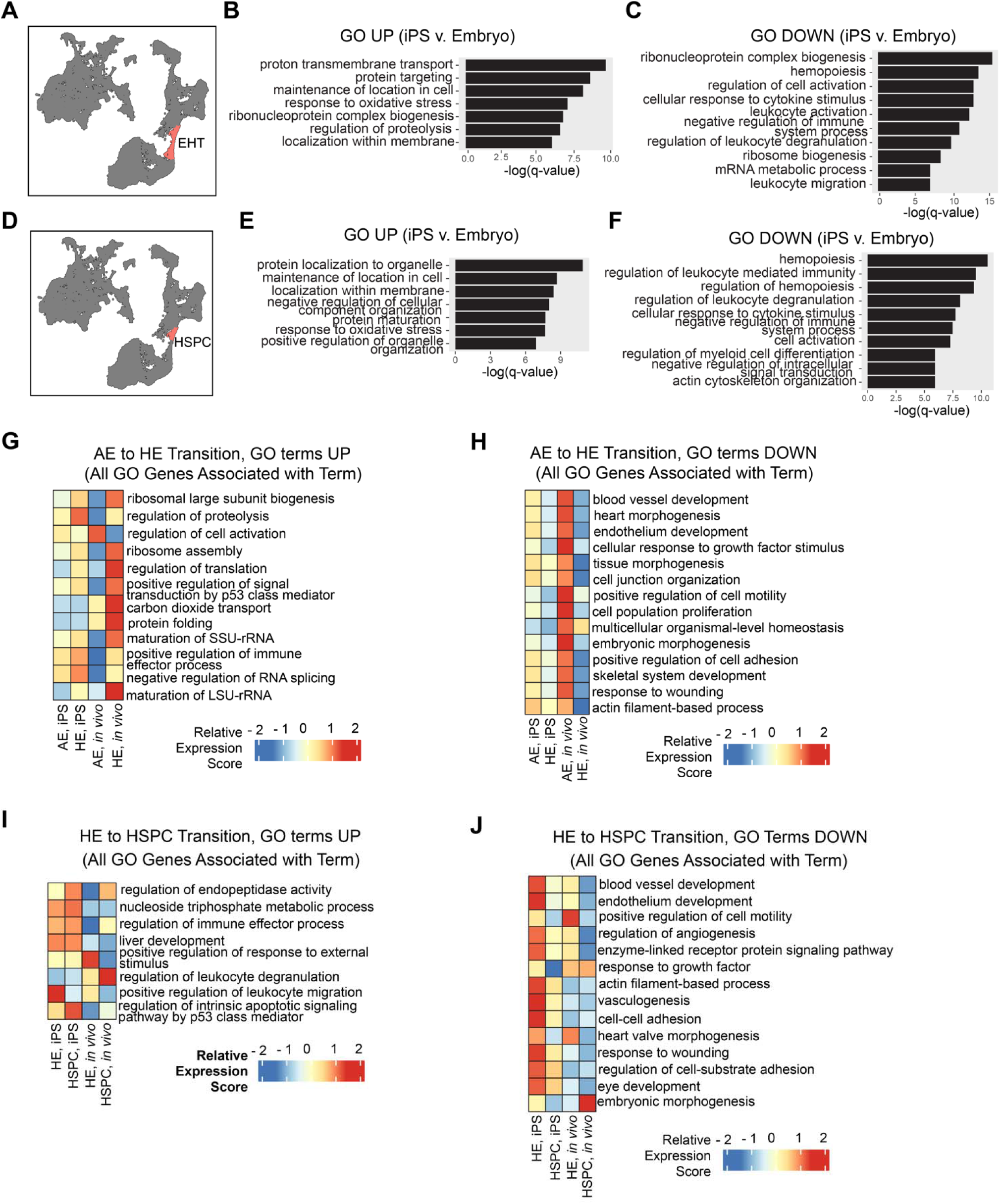
Expanded comparative molecular analysis of EHT *in vitro* and *in vivo*. Related to Figure 4. (**A**) Combined HE (cluster 75) and early HSPC (cluster 74) population undergoing EHT used for differential gene expression analysis. (**B,C**) Gene ontology (GO) terms significantly upregulated (B) or downregulated (C) in iPSC-derived vs. HSC-competent human embryo EHT (Metascape, q < 0.05). (**D**) Early HSPC (cluster 74) population used for differential gene expression analysis. (**E,F**) GO terms significantly upregulated (E) or downregulated (F) in iPSC-derived vs. HSC-competent human embryo early HSPC (Metascape, q < 0.05). (**G**) Relative aggregate expression score of all genes associated with the GO terms upregulated in the AE to HE transition in Figure 4E. (**H**) Relative aggregate expression score of all genes associated with the GO terms downregulated in the AE to HE transition in Figure 4F. (**I**) Relative aggregate expression score of all genes associated with the GO terms upregulated in the HE to HSPC transition in Figure 4H. (**J**) Relative aggregate expression score of all genes associated with the GO terms downregulated in the HE to HSPC transition in Figure 4I. For (G-J), aggregate gene expression scores were calculated and combined in scaled matrices which were used for heatmap generation.

**Figure S5.**
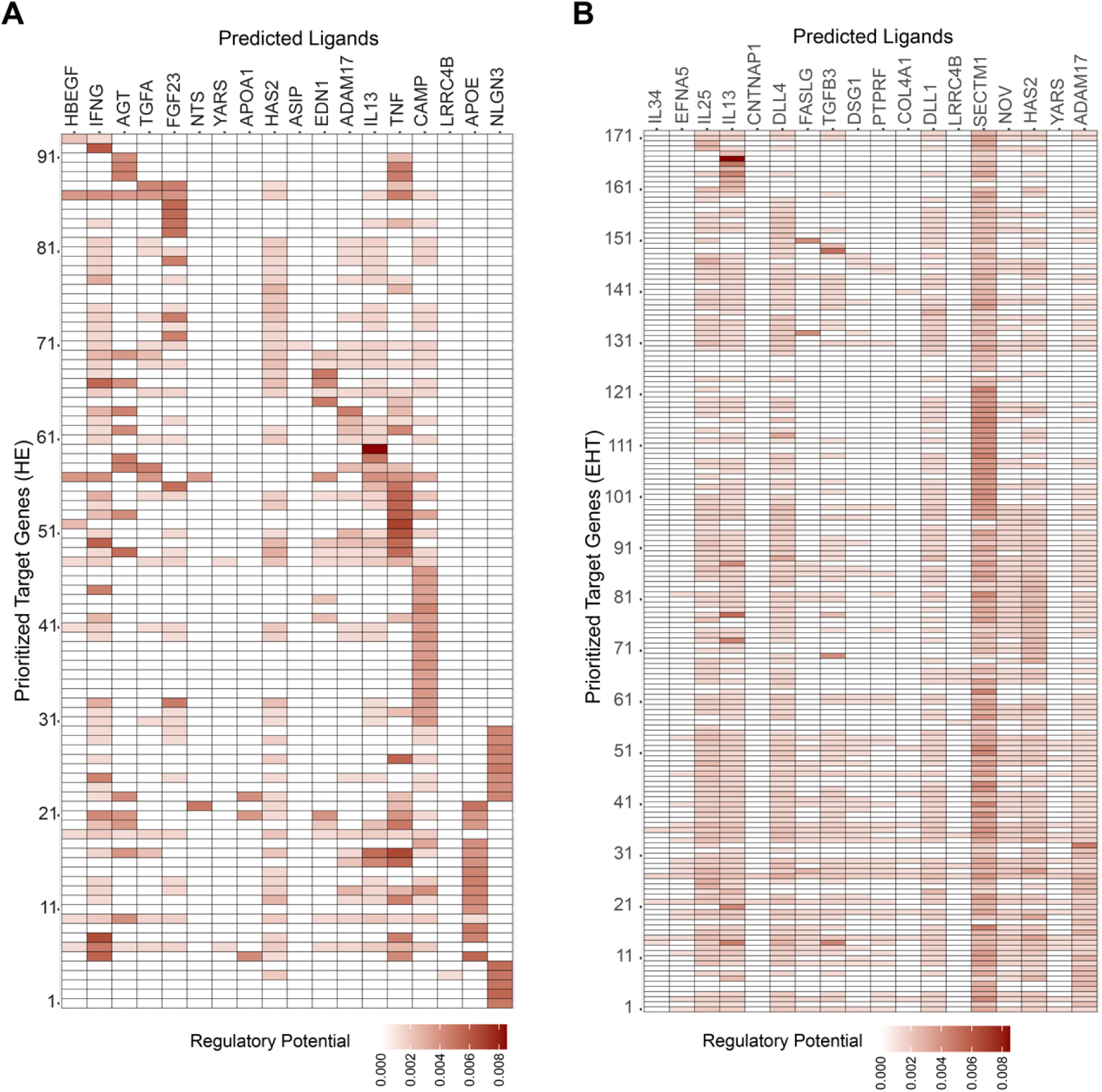
Expanded map of ligand-receptor interactions to inform iPSC differentiation. Related to Figure 5. (**A**) Regulatory potential of each predicted ligand for HE (columns) to regulate target DEGs (row); DEGs are listed in Table S5. (**B**) Regulatory potential of each predicted ligand for EHT (HE + HSPC) to regulate target DEGs; DEGs are listed in Table S5.

**Figure S6.**
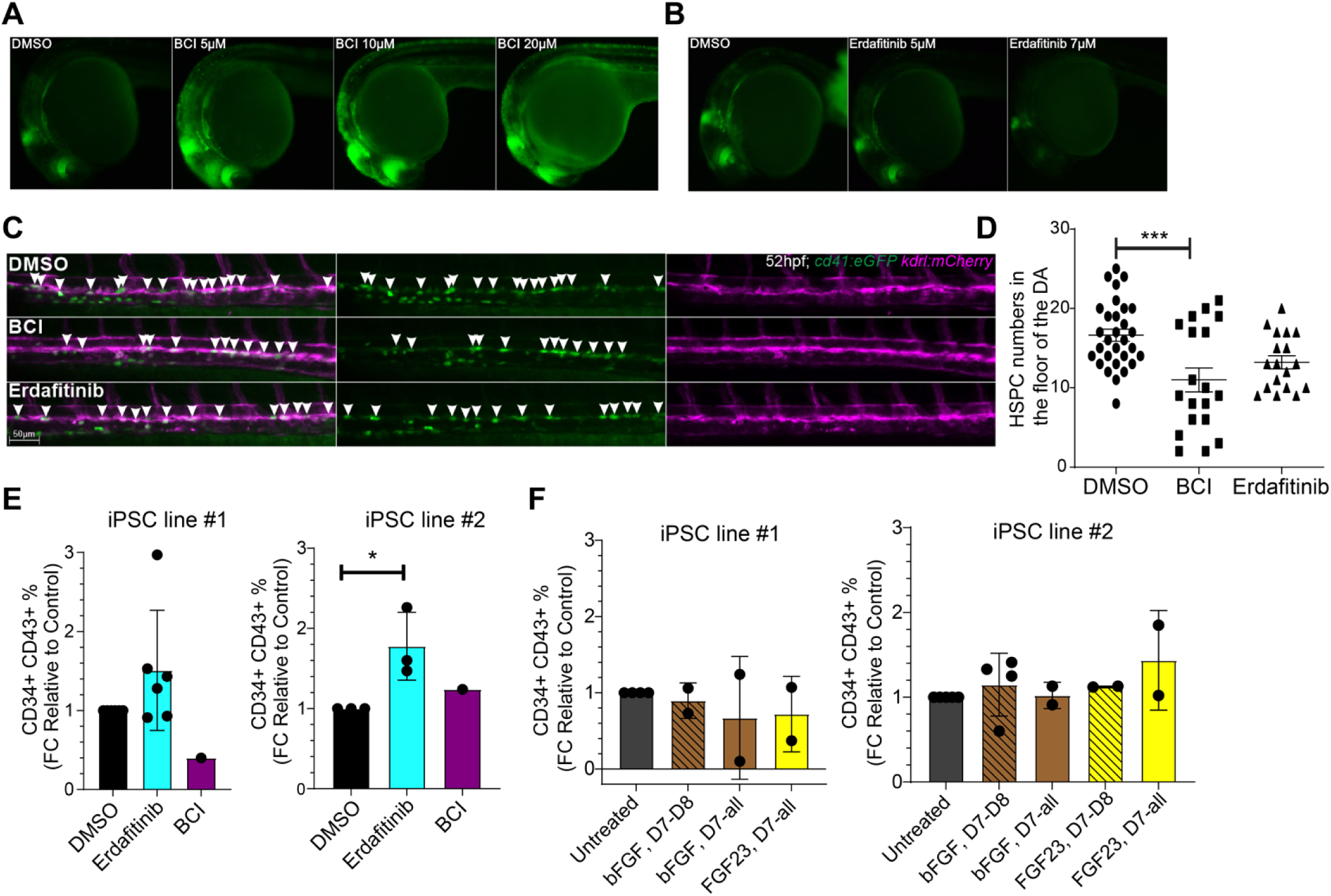
Expanded chemical modulation of FGF signaling during EHT regulates hematopoiesis *in vivo* and *in vitro*. Related to Figure 6. (**A**) Titration of BCI concentration in zebrafish using the *dusp6:d2EGFP* FGF reporter. 10 µM was used in experiments in Figure 6. (**B**) Titration of erdafitinib in zebrafish using the *dusp6:d2EGFP* FGF reporter; 7 µM was used in experiments in Figure 6. (**C-D**) Representative images (C) and quantification (D) of the number of HSPCs (white arrows) in the DA following treatment with an FGF small molecule-inhibitor (erdafitinib) or agonist (BCI) prior to EHT. Zebrafish were treated with 7 µM erdafitinib, 10 µM BCI, or vehicle (DMSO) 16 - 24 hpf and *kdrl:mCherry^+^; cd41:eGFP^+^* HSPCs were quantified at 52 hpf. Mean ± s.e.m. of 30 embryos in 2 independent experiments, unpaired two-tailed t-test (*p<0.05, ***p<0.001). (**E**) Percent of CD34^+^CD43^+^ iPSC-derived HSPCs following treatment with 0.1 µM erdafitinib or 10 µM BCI as shown in Figure 6E for the two different iPSC lines (lines 1 and 2). Mean ± s.d. (single outlier point removed), paired two-tailed Student’s t-test (*p<0.05). (**F**) Percent of CD34^+^CD43^+^ iPSC-derived HSPCs following treatment with FGF ligands (bFGF, FGF23) as shown in Figure 6F for the two different iPSC lines (lines 1 and 2).

## Notes

https://cellxgene.cziscience.com/collections/4a2c25af-558a-45fc-bc9a-54ec44a1d63f

### Summary of Updates

Layout and style of figures updated to improve data presentation; panels moved between main and supplemental figures; supplemental figures moved to appear at the end; competing interests updated. This version also includes the link to an interactive online interface for exploring the data.

